# Implicit adaptation’s effect on sensorimotor and motor confidence

**DOI:** 10.64898/2025.12.15.694412

**Authors:** Marissa H. Evans, Jordan A. Taylor, Michael S. Landy

**Affiliations:** Dept. of Psychology, New York University, New York, NY, USA; Dept. of Psychology, Princeton University, Princeton, NJ, USA; Princeton Neuroscience Institute, Princeton University, Princeton NJ, USA; Center for Neural Science, New York University, New York, NY, USA

**Keywords:** implicit, adaptation, reaching, wager, confidence, motor, metacognition

## Abstract

Sensorimotor adaptation maintains movement accuracy by counteracting perturbations or miscalibrations. It can operate explicitly, by consciously adjusting motor plans to correct errors, or implicitly, by automatically recalibrating sensorimotor mappings without altering the motor plan. While explicit adaptation is known to reduce sensorimotor confidence—the perceived likelihood of successful action—it is unclear whether implicit adaptation similarly affects confidence in sensorimotor judgments or motor awareness (knowledge of one’s own limb position). To investigate this, participants made reaching movements to visual targets without seeing their hand. Cursor feedback followed an “error-clamped” trajectory: its radial position matched the hand, but its angular direction was fixed and independent of actual hand direction, a manipulation participants were told to ignore. The clamp direction varied sinusoidally over trials (±10°; 12 cycles per session). Participants reported confidence by adjusting the size of an arc centered on the target or, in another task, centered on reported reach direction; larger arcs indicated lower confidence. Points were awarded when the arc encompassed the true reach direction, with fewer points for larger arcs, encouraging accurate and meaningful confidence reports. Fourier analysis of reach and report time series revealed a strong 12-cycle component in both, demonstrating robust implicit adaptation and corresponding changes in motor awareness. These findings indicate that although implicit adaptation operates unconsciously, the resulting mismatch between motor plans and proprioceptive signals can bias judgments of reach outcomes. However, confidence judgments were not consistently affected, suggesting that sensorimotor confidence and confidence in proprioceptive awareness may rely on partially distinct mechanisms.

**New & Noteworthy:** This work is among the first to look at how implicit adaptation influences meta-judgments regarding motor execution and internal states. It identifies a dissociation between sensorimotor confidence and motor-awareness confidence.

## INTRODUCTION

When using our bodies to act on the world around us, we often have to evaluate the internal state of our body and judge the accuracy of our motor execution. The former has been defined as motor awareness, which is important when executing an action without an external goal, such as making a hand gesture or performing a dance move (1, 2). It often depends wholly on your sense of proprioception (sense of the body’s position in space) and kinesthetics (sense of the body’s movement and pose) (3). Confidence in this motor-awareness judgment reflects an assessment of uncertainty in internal sensory systems (4). In contrast, sensorimotor confidence regards the assessment of the outward success of an action, which often is sensory directed and has a perceptual– or task-level goal (5, 6). Examples include throwing a dart, reaching for a glass, or catching a ball ⏤ any action where success or failure is determined by the action’s outcome relative to a goal (7).

When a discrepancy is detected between an action’s outcome and intended goal, the motor system adapts the action — either consciously or unconsciously — to reduce the mismatch (8). Such adjustments occur frequently in daily life, whether it be adapting to a new pair of eyeglasses, adjusting grip force when lifting an unexpectedly heavy cup, or when muscles become fatigued. More deliberate corrections can also be made to combat a consciously sensed exterior force, such as adjusting one’s throw to account for strong winds. Motor adaptation is typically studied using reach-perturbation paradigms such as visuomotor rotations or force-field adaptation, in which participants readily adjust their movement direction or force output to counteract the imposed perturbations. Whether these adjustments occur automatically (implicitly) or deliberately (explicit) results in different outcomes, each with its own unique features, and are often dependent on task conditions, such as type of feedback or strength of the perturbation (9, 10). Although the mechanisms of motor adaptation have been the focus of a wide body of research (9, 11–14), there has been little attention to how it influences sensorimotor or motor-awareness confidence. This is of particular importance because adaptation demands accurate and confident assessments of internal states to guide recalibration for the success of future movements (10, 15–17).

Experimentally, motor adaptation can be induced in a reaching task by perturbing the visual feedback shown to the observer, with the strongest adaptation stemming from full trajectory feedback (16, 18–21). In the case of explicit adaptation the participant has control of the perturbed feedback, so they make explicit changes to the motor plan to shift it towards the perceived goal (22, 23). During implicit adaptation the motor plan remains the same, but unconscious shifts are made to reduce potential mistakes (9, 16, 24–26). The perceived errors must be tied to the motor plan and relevant in the visual scene (17, 27). Importantly, the shifts caused by implicit adaptation are not intentional (22); however, they may still be sensed by the observer if proprioceptive input is interpreted in the same polar coordinates (28–33).

Previously, we examined how explicit adaptation affected sensorimotor confidence, finding that confidence is reduced sharply over the course of adaptation to a newly perturbed environment (34). Confidence does not appear to stabilize until several trials after the adaptation has plateaued, and may not always return to pre-perturbation levels. This mirrors findings in the sensorimotor-learning literature showing that perceived uncertainty and variability in outcomes during adaptation can impair confidence even when motor performance is improving (35, 36).

In the present work, we turn our attention to implicit adaptation, asking how it affects sensorimotor and motor-awareness confidence. To induce implicit adaptation, in relative isolation of explicit strategies (37, 38), we leveraged the task-irrelevant error-clamp feedback paradigm (9, 16). In this paradigm, participants were provided with online cursor feedback whose radial position was yoked to their movement velocity, but whose angular direction was fixed relative to the target. Participants were explicitly informed that the clamped feedback was not influenced by their actual movement direction and were instructed to ignore it and reach directly to the target. On different trials, participants were queried about their sensorimotor or motor-awareness confidence. For sensorimotor-confidence trials, participants rated how confident they were that they hit the target, consistent with paradigms that assess outcome-based metacognition (6, 15, 36). For motor-awareness confidence trials, participants first reported their perceived hand position and then rated their confidence in the accuracy of that perception, aligning with frameworks that investigate the supra-modal mechanisms of metacognition (39, 40).

To ensure that participants were required to judge their momentary sensorimotor and motor awareness confidence, we induced trial-by-trial variance in the error-clamped feedback such that it followed a subtle, periodic function. Beyond requiring continuous adjustment in meta-level cognitive judgement, this approach also offers an analytic advantage from a system-identification perspective: the response function of implicit adaptation, sensorimotor confidence, and motor-awareness confidence can be evaluated relative to periodicity of the experimentally controlled time-varying perturbation (31, 41, 42). What’s more, the effect of implicit adaptation could be small — if present — and this approach should enhance detection sensitivity of subtle changes in adaptation and confidence ( 2, 37).

We found that despite knowledge of the task-irrelevant perturbation, all participants consistently exhibited gradual deviations in reach direction away from the target, in a direction opposite to the clamped feedback, which is viewed as a hallmark of unconscious, implicit adaptation driven by sensory-prediction errors (20, 38). Importantly, reach adaptation followed a periodic function in response to our imposed sinusoidal perturbation, enabling the use of Fourier analysis to detect frequency components matching those embedded in the perturbation. Fourier analysis is highly sensitive to periodic structure and can uncover subtle fluctuations that might be missed by traditional correlation-based approaches (37).

Of more interest, the sinusoidal perturbations enabled applying Fourier analysis to participants’ confidence judgments. Participants performed two confidence tasks while undergoing implicit adaptation, one that measured sensorimotor confidence and another that measured motor-awareness confidence. We found that implicit adaptation affected confidence for only half of our participants: confidence dropped on trials where the perturbation resulted in a larger mismatch between the hand and cursor than was expected. For the other half, confidence did not fluctuate in phase with changes to the visual feedback or adapted hand position.This suggests that although every participant successfully exhibited implicit adaptation, some did not use the shifted proprioceptive hand position or the error-clamped visual feedback when forming their confidence judgments. When confidence fluctuated with the error-clamp feedback it decreased whenever the visual feedback conflicted with the actual reach direction. For the sensorimotor confidence task confidence should change with the adaptation magnitude to maximize expected gain, however for the motor-awareness confidence task the report should reflect the adaptation, with confidence untethered to report direction. However we do see a fluctuation of confidence coinciding with the adaptation, indicating that confidence is influenced by visual feedback and proprioceptive mismatch, even when not task relevant.

## METHODS

### Ethics Statement

The study’s experimental design and recruitment process received approval from the New York University Committee on Activities Involving Human Subjects. All participants provided informed written consent and received financial compensation for their time.

### Participants

Twenty individuals were selected from the New York University student body (average age: 27.3 years, SD: 4.8 years, 10 males, one left-handed). None of the participants were familiar with the experimental design. All participants had either normal or corrected-to-normal vision, no restrictions on their right arm’s mobility, and reported no motor abnormalities. All participants completed both the motor-awareness and sensorimotor tasks across two sessions.

### Apparatus

Participants were seated at a desk facing a custom-made support frame that held a mirror and a frosted-glass screen above the table’s surface (Fig. 1). A projector (Hitachi CP-X3010) mounted above the screen facing downward displayed visual stimuli on the glass screen, which was positioned above the mirror. The mirror was placed equidistant between the screen and an input tablet device (Cintiq 22; Wacom, Vancouver, WA) lying flat on the desk. The mirror reflected the underside of the screen (showing the display) back toward the participant, creating the perceptual experience that the stimuli were on the same spatial plane as the tablet surface. Participants were positioned in a chin rest, with their line of sight above the mirror but below the screen, and were unable to see their hands during the task. The stimuli were presented against a mid-gray background. Reach end points were recorded using a stylus, held in the right hand, on the digitizing tablet at a sampling rate of 200 Hz. The experimental software was custom-built in Matlab (Mathworks, Natick, MA, USA), utilizing the PsychToolBox extension (43–45), and ran on a Dell Optiplex 9020 PC operating on Windows 10 (Microsoft, Redmond, WA, USA).

**Fig 1.**
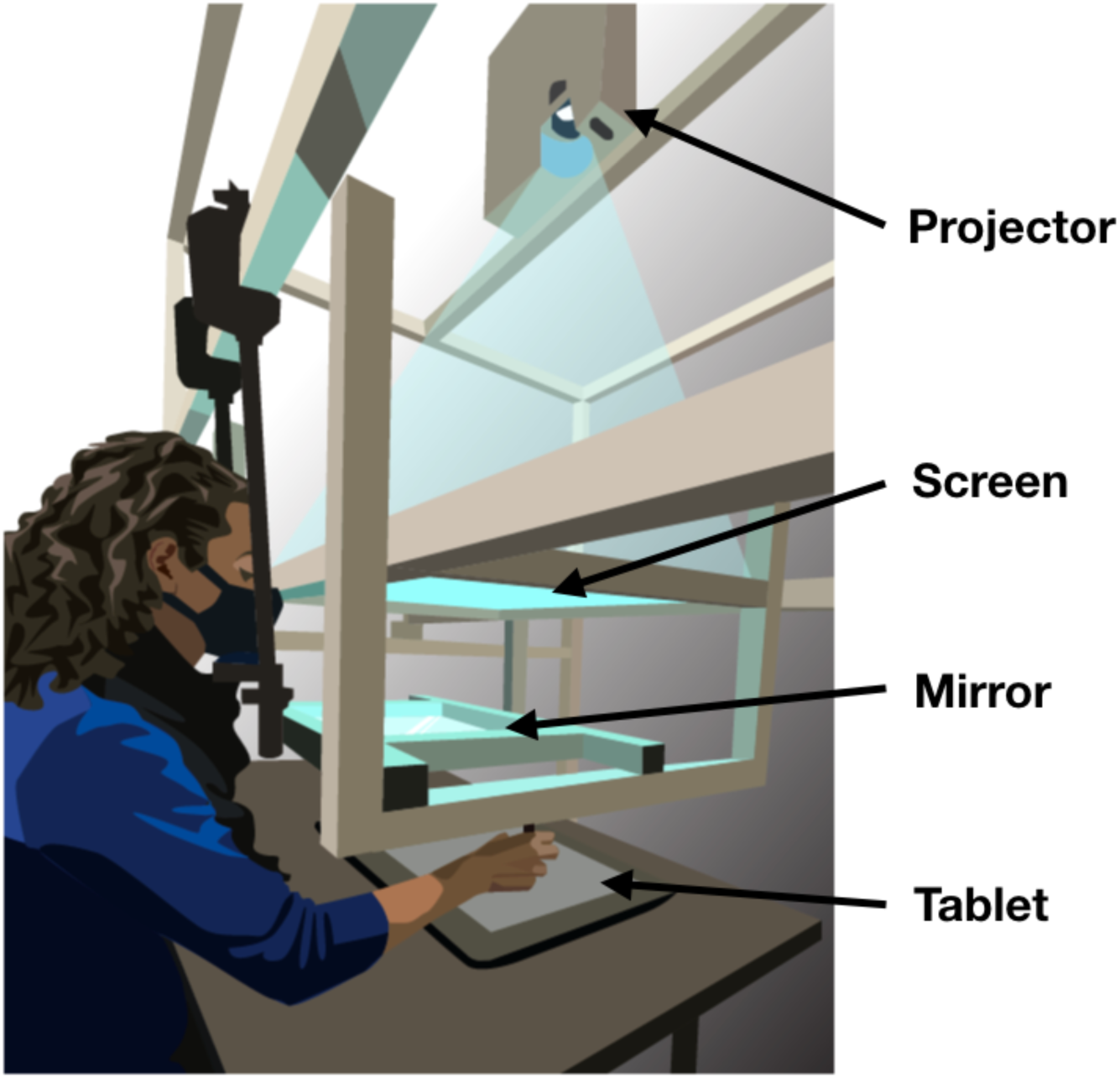
Apparatus. Participants responded with reach movements recorded by a digitizing tablet placed on a table. A mirror was suspended between a frosted-glass screen and the tablet surface in the same horizontal plane. The projector above the glass screen projected stimuli downward. Participants viewed the mirror from a fixed head position and could not see the tablet or their hand while performing reaches with a stylus. The virtual image of the display was in the plane of the tablet. Figure from Fassold, Locke & Landy, (2023) Plos Comput Biol. 19(6):e1010740.

Before each session, the tablet was aligned with the on-screen display by using a partially transparent half-silvered mirror in place of the fully silvered mirror, allowing both the display and the hand to be visible. Participants touched the tablet with the stylus at each point in a 3×3 grid of dots across the projected screen area for calibration. The resulting data were used to calculate an affine transformation between screen and tablet coordinates for each session for every participant using least-squares estimation. The calibration data from each session were only used to convert projected stimuli and recorded reach end points within that same session. Additionally, participants used a Griffin PowerMate control knob with their left hand for responses during both tasks.

### Motor-Awareness Pre-Test

The first twenty trials of both sessions were practice trials with online, veridical cursor feedback of the reach. This allowed participants to get used to reaching without seeing their hand and to familiarize themselves with the apparatus. For the next 40 trials, participants were asked to report their perceived reach direction without any feedback. These data afforded independent assessments of proprioceptive and motor uncertainty outside the main task. The same behavioral setup and model fit were used as in previous research from the lab (34).

Previous studies have demonstrated individual differences in proprioception (6, 26), primarily operationalized as one’s accuracy in localizing the position of an unseen hand. Thus, we sought to assess the participant’s proprioceptive uncertainty (motor awareness) separately from the confidence tasks to provide a value for each participant prior to adaptation. To achieve this, participants were asked to report the reach direction of their unseen reaches, and a Bayesian-inference model was applied to these reports. This control task was conducted before the main confidence task trials of each session and included 40 experimental trials, for a total of 80 trials used in the analysis.

During each trial, participants positioned the stylus within a 7 mm annulus at the bottom of the screen (Fig. 2). When the stylus was within 2 cm of the annulus center, visual feedback of the stylus location (a 4.5 mm diameter white dot) was provided. Once the cursor was inside the annulus, a target (a 4.5 mm white dot) appeared 150 mm from the start-location annulus, directly in front of the participant. After 800 ms, the target turned green, indicating to the participant to initiate a slicing reach through the target location. Upon passing the target distance (150mm), the reach was completed and participants were instructed to keep the right hand in place at the reach end point to encourage the use of ongoing proprioceptive signals while they used their left hand to report their perceived reach direction. This was accomplished by rotating a dial, which moved a visual cursor along an invisible arc (starting direction randomized), centered on the start point, intersecting the target (radius 150 mm from the starting annulus). The target location remained constant throughout all trials, and was visible during the direction report. No feedback was provided during this task. If participants (1) missed the response window (800 ms to start movement following the go cue and 1200 ms to complete the reach after reach initiation), (2) moved their hand out of the annulus before the go cue, or (3) lifted their hand off the tablet surface during the reach the they received a visual warning asking them to repeat their response.

**Fig 2:**
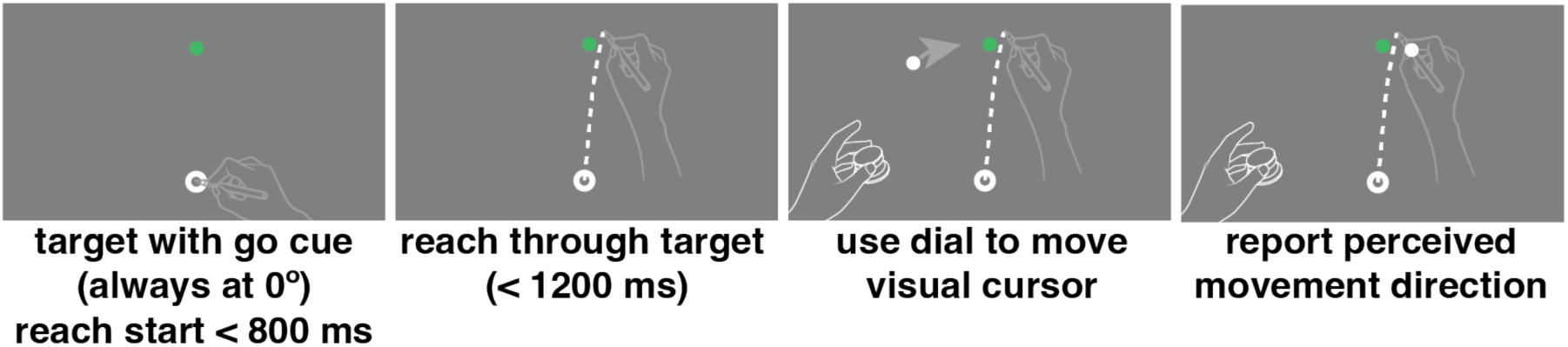
Motor-awareness task trial sequence. Participants began the trial with their hand positioned in the starting annulus at the bottom of the screen. Once the target turned green they performed a slicing reach through the target with no visual feedback of hand position. Leaving the right hand in position at the end of the reach they used the left hand to rotate a dial that moved a white dot on the screen in an arc 150 mm from the starting point. Once the visual cursor was positioned at their perceived reach direction they pressed down on the knob to save the report and complete the trial.

### Motor-Awareness Model

We modeled the data from the motor-awareness task separately from the data from the main confidence tasks (described in the next section) in order to obtain independent estimates of proprioceptive uncertainty for each participant. We use the same model as we used in past work in our lab to fit motor and proprioceptive error in this task (6, 34). Following our previous work, we assumed an ideal-performance model that combines prior knowledge of typical reach directions and the noisy sensed reach direction. The model determines the maximum a posteriori (MAP) estimate of the true reach direction given that the reach objective was to slice through the target. The participant aimed at the target direction, straight ahead (i.e., 0°) in all trials. We assume that the reach is noisy and unbiased and the distribution of reach directions has motor variance, 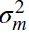, that is known to the participant. That is, the prior distribution, *p*(*e*), of the reach direction, *e*, is 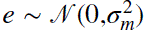. After the reach, the participant has a noisy proprioceptive measurement, *s*, of movement direction, with proprioceptive variance, *σ_p_*, that is also known to the participant. Thus, 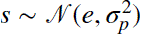. The corresponding motor and proprioceptive reliabilities are 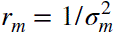 and 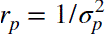, respectively. The participant knows *s* and tries to estimate *e*. We assume that the variance of the visual estimate of target direction is negligible relative to motor and proprioceptive noise and thus treat target direction as known precisely. From the participant’s perspective, the likelihood function is then 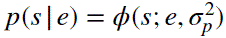, where *ϕ* denotes the value of a normal distribution at the sensed direction, *s*, with a mean of the true reach direction, (unknown to the participant), and variance 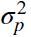. From the participant’s perspective, the posterior distribution is 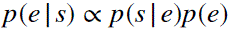, and the participant estimates the end point location as the maximum a posteriori (MAP) estimate of reach direction, and thus indicates direction:

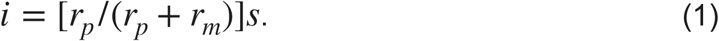

In this model motor noise, *σ_m_*, and proprioceptive noise, *σ_p_*, are the parameters we fit to the data. In this task we assume there is no uncertainty in reporting the perceived reach direction, that is, we do not include additional noise from using the dial to indicate the direction in the model. We estimated these parameters by maximum likelihood. Thus, from the experimenter’s perspective, the likelihood of the data, *d*, from a single trial *j*, where 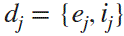, is

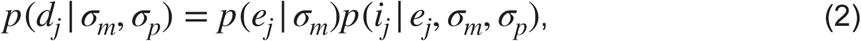

where the likelihood of the end point is 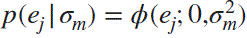 and the likelihood of the reported end point, *i_j_*, is

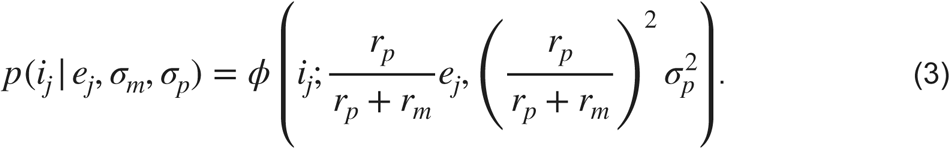

Our estimates of a participant’s motor and proprioceptive noise are the values of *σ_m_* and *σ_p_* that maximize the overall likelihood, or equivalently the log likelihood, across all trials, 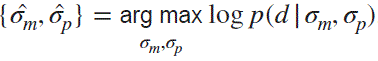. The log likelihood across all trials can be summed based on our assumption of independence.

### Sensorimotor-confidence task

During the sensorimotor-confidence task the participant placed the right hand in the starting annulus, following which a target (4 mm white dot) appeared 150 mm in front of the participant (0°; straight ahead of the observer and at the same location every trial). After 800 ms the target turned green, signaling that the reach should be executed (Fig. 3). While visual feedback of a cursor (4.5 mm dot) was shown to the observer following each reach, a sinusoidal error clamp was applied to the angular direction of the cursor feedback (1 cycle every 20 trials) that was fully controlled by the experimenter. The radial position of the cursor faithfully followed the radial movement of the hand. The underlying sinusoid controlling the direction of the error-clamp feedback had a maximum amplitude of +/-10°, starting at 0° for the first trial. The participant was made aware of the clamped feedback and was instructed to ignore it, attempting to only reach directly to the target with their hand as accurately as possible. Just as in the primary motor-awareness task, after completion of the reach the participants were instructed to keep their right hand in place (at its terminal position) while using their left hand to operate a control knob to make the confidence judgment. The confidence judgment was operationalized as an arc intersecting the target location with a radius of 150 mm from the starting annulus. The arc extended in both directions equidistantly away from the target, positioned at 0° straight ahead of the participant. The initial arc’s subtense was randomly selected on each trial from a uniform distribution ranging from 1 to 40°. Small arcs equated to high confidence (“I’m confident I hit the target.”) and large arcs meant low confidence (“I’m not confident I hit the target.”). Participants received points if their confidence arc enclosed the true reach direction. If the confidence arc did not include the reach direction, zero points were awarded on that trial. If the arc did include the reach direction, the points associated with an arc of that size were awarded. The reward was higher for smaller arc sizes, decreasing linearly from 10 points for arcs of size 2° or less to 0 points for 40° or more. Participants were briefed on the relationship between reach direction, arc size, and points, and instructed to earn as many points as possible within these parameters. This reward structure emphasizes accurate performance (greater reward for smaller arc size) while also fostering a reasonable confidence report based on performance in each trial (6), a technique previously successful for measuring spatial-memory confidence (46), and aligns the confidence report with the error scale, allowing for direct comparison between error magnitude and confidence. Every 60 trials participants were shown a leaderboard displaying all previous participants’ scores for the previous 60 trials, motivating participants to maximize their expected gain. The leaderboard only displayed the order of participants’ scores as first place, second place, etc. If the participant did not score higher than any of the other participants, then they were always shown in the 6th place on the board to prevent them from losing motivation. Since the adaptation in this task was implicit, we did not want to give participants an idea of how their behavior was affected by adaptation, as it could in turn change how they reported confidence.

**Fig 3:**
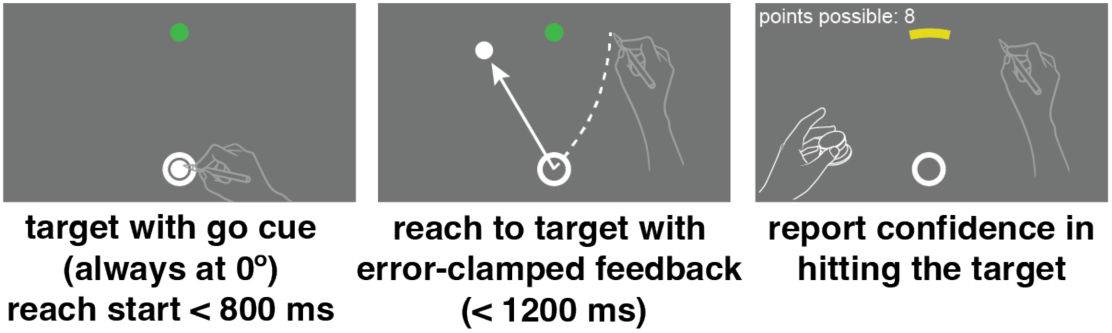
Sensorimotor-confidence task trial sequence. Participants started the trial with their hand positioned in the starting annulus at the bottom of the screen. Once the target turned green they performed a slicing reach through the target with error-clamped visual feedback. Leaving their right hand in position at the end of the reach they used the left hand to rotate a dial that set the width of an arc centered at the target location to indicate their confidence in whether they hit the target. Large arcs meant low confidence and fewer points, and small arcs high confidence. There was a points wager associated with the confidence judgment, where all points possible were awarded only if the true reach direction was enclosed in the arc.

Since the confidence arc appeared at a random width it was verified that participants were actively participating in the task by a non-significant correlation between these starting positions and the final reports. One participant was removed from the task because they did not change the confidence-arc sizes after they appeared and was subsequently excluded from any analyses. Trials were aborted if the participant missed the response window, moved the hand from the start location before the go cue, or lifted the hand off the tablet during the reach. In such cases the trials were repeated until successful.

### Motor-awareness confidence task

In the other session of our task we asked participants to perform a motor-awareness confidence judgment instead of a sensorimotor confidence judgment. This is an important distinction because in this task the underlying question is “how well do you know where your hand is?” While in the previous sensorimotor confidence task the question was “how well did you hit the target?”. Sensorimotor confidence is specific to the external goal of the task, while motor-awareness confidence is about your knowledge of your body’s position, regardless of the vicinity to the target. In the motor-awareness confidence task the participant’s behavior is nearly identical to the sensorimotor task, however now they make an additional report of their perceived hand position, then rate their confidence in that report. The observer made a slicing reach through the target while being shown visual feedback controlled by the experimenter that sinusoidally shifted direction with a cycle of 20 trials and a maximum amplitude of 10° away from the target. The feedback moved at the same speed as the participant but the direction of the feedback was fixed on a given trial and not affected by any changes the participant made to their movement direction. Just as in the primary motor-awareness task, after reach completion the participant kept the right hand fixed at the end point of the reach. Using the left hand, they moved a white dot along an invisible arc (radius 150 mm from the starting annulus) with its apex centered at the target location using a control knob to report perceived hand direction.

Once the perceived-direction report was complete they then made a second report, a confidence judgment about whether their report was close to their true reach direction. For the confidence report, participants adjusted the size of an arc (radius: 150 mm from the starting annulus) centered on the reported location using a control knob. The arc’s magnitude changed uniformly in both directions away from the reported location. The initial arc’s subtense was randomly selected on each trial from a uniform distribution ranging from 1° to 20°. Small arcs equated to high confidence (“I’m confident my reported location is accurate.”) and large arcs meant low confidence (“I’m not confident my reported location is accurate.”). Participants once again received points if their confidence arc enclosed the true reach direction. If the confidence arc did not include the reach direction, zero points were awarded on that trial. If the arc did include the reach direction, the points associated with an arc of that size were awarded. The reward was higher for smaller arc sizes, decreasing linearly from 10 points for arcs of size 2° or less to 0 points for 20° or more. All other aspects of the trial were identical to the sensorimotor-confidence task.

### Permutation testing of Fourier analysis

The error-clamped visual feedback completed a full cycle every 20 trials, for a total of 12 cycles per session for both tasks. If the error-clamped feedback induced adaptation, we would expect a Fourier component of 12 cycles per session for the reach direction. Likewise, if participants maintain an accurate representation of their sensed hand position, then we would also expect a Fourier component of 12 cycles per session. However, for the confidence report, we may expect a 24 cycle per session Fourier component because confidence could be reduced whenever the error-clamp display or the reach direction diverges from the target, which occurs twice per error-clamp cycle (in the clockwise and counter-clockwise directions).

To determine whether the amplitude of the expected frequency component was statistically significant we used a permutation test. First a Fourier analysis of the full session was run to identify the amplitude of the 12 or 24 cycles per session component. Then, the trial order was shuffled, disrupting any pattern present in the sequence. The Fourier analysis was run on this shuffled data set and this process was repeated 100,000 times. If the Fourier component from the baseline analysis had an amplitude larger than 99% of the shuffled sessions, that Fourier component was considered significant. If a frequency was significant, the phase (trial lag) and amplitude were considered in further analyses.

## RESULTS

### Motor Awareness (pre-test)

The goal of the motor-awareness task was to provide estimates of each participant’s motor and proprioceptive uncertainty, *σ_m_* and *σ_p_*, that were independent of the main confidence tasks. In this motor-awareness task participants repeatedly made ballistic reaches to a fixed target location, and then reported the perceived end point direction. An identical behavioral design was used by Evans & Landy (34); all aspects of the task remained the same and the data were analyzed in the same way. Parameter values for each participant were determined using maximum-likelihood estimation.

Across the 20 participants the average *σ_m_* estimate was 3.99° (SD: 1.37°) and the average *σ_p_* estimate was 7.16° (SD: 3.65°; Fig. 5A). While motor error was similar for all participants, there was a wider spread of proprioceptive uncertainty, as also found in previous research (6, 34). We found that the reliabilities from the model fit aligned with the data for each participant (Fig. 5B).

**Fig 4:**
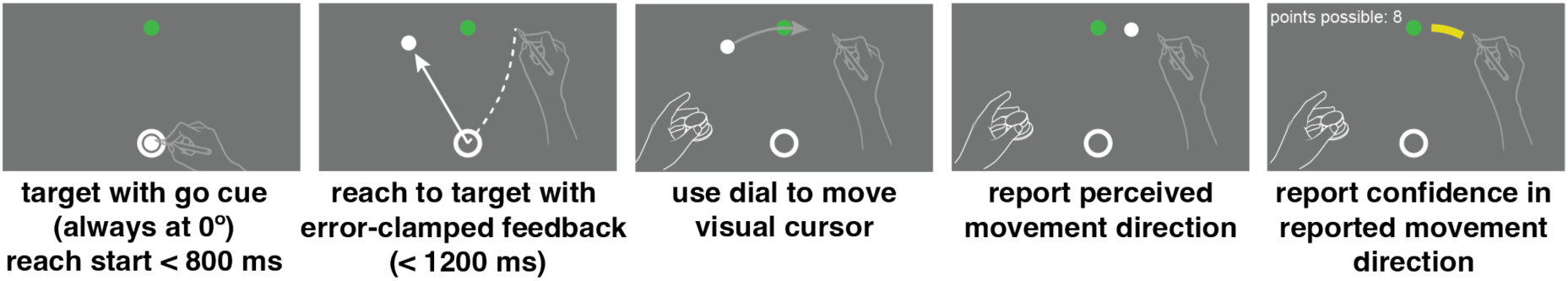
Motor-awareness confidence task trial sequence. Participants started the trial with their hand positioned in the starting annulus at the bottom of the screen. Once the target turned green they performed a slicing reach through the target with error-clamped visual feedback. The direction of the error clamp varied sinusoidally over trials. Leaving the right hand in position at the end of the reach, they used the left hand to rotate a dial that moved a white dot on the screen in an arc always 150 mm from the starting point. Once the dot was positioned at their perceived reach direction they pressed down on the knob to save the report. They then used the dial a second time to set the width of an arc centered at their reported direction to indicate their confidence in their direction report. Large arcs meant low confidence and fewer points, and small arcs high confidence. There was a points wager associated with the confidence judgment, where all points possible were awarded only if the true reach direction was enclosed within the arc.

**Fig 5:**
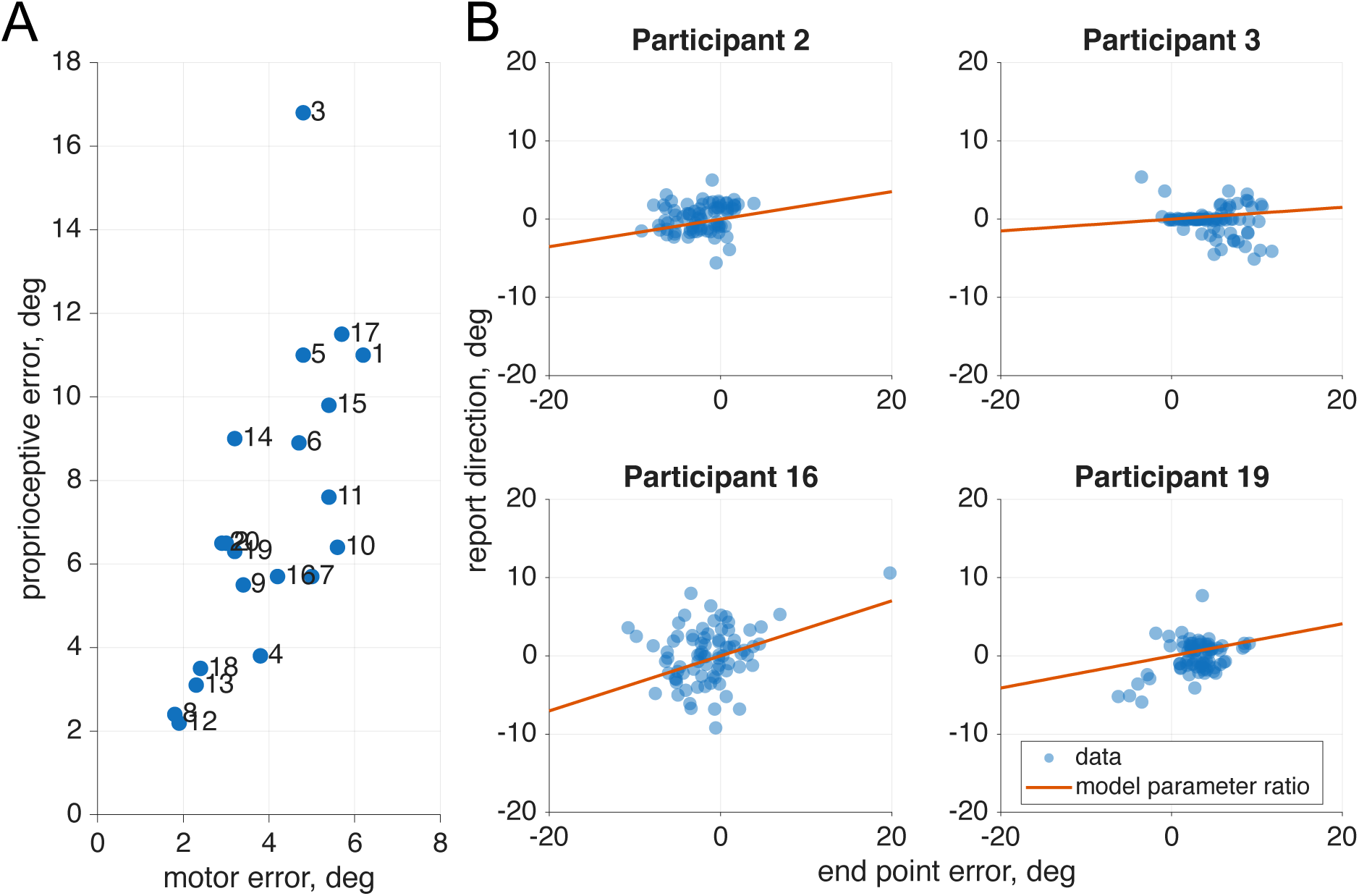
Motor and proprioceptive uncertainty from the motor-awareness task. A. Proprioceptive and motor uncertainty were separately fit from 40 trials after the practice from the start of both sessions-for a total of 80 trials used in the fit, in which the participant made motor-awareness judgments. Consistent with previous research, there is a larger variation in proprioceptive uncertainty across participants than there is in motor uncertainty. B. For four sample participants (used as examples throughout the paper, selected based on significance of phase components in the main confidence task) the end point error and reported reach direction were plotted and overlaid with a line following the slope of the reliabilities of the parameters fit by the model. We can see that our model captures trends within the data. Plots for all participants are in Supplement 1.

### Sensorimotor Confidence

Sensorimotor and motor-awareness confidence are both dependent on multiple sensory inputs (e.g., proprioception, vision) and prior information (15, 36, 47). As shown in previous research (6), during a sensorimotor confidence task, inclusion of proprioceptive input makes confidence judgments closer to optimal, particularly when incentives are used (48). However, participants can still perform well without proprioceptive input when prior experience is reliable and unperturbed (49), consistent with Bayesian models of sensory integration (4). Once a perturbation is introduced, however, trial-specific information (e.g., proprioception), becomes increasingly valuable as the reliability of prior visual feedback is reduced (26, 29, 50). In contrast, motor-awareness confidence is thought to rely primarily on proprioception and its associated uncertainty (51), as there is no external goal to verify the correctness of the action. Rather, only internal sensory estimates are available to judge movement outcome (40). During our implicit-adaptation task using a sinusoidal error-clamp paradigm, the participant has access to several information sources on each trial: the target location, the internal motor plan, the visual feedback direction (known to be uninformative), and their sensed hand location. Because sensorimotor confidence is related to the success of an action in arriving at the target, we expect it to decrease when hand position deviates from the target, and increase when the hand is closer, consistent with performance monitoring (36, 52). In contrast, motor-awareness confidence is based on a self-generated direction judgment, and thus should not be influenced by the distance of the hand or the end point feedback from the target (i.e., target error). If we do see a modulation in confidence correlated with the distance the reported direction is from the target, then we conclude that visual feedback modulates confidence indirectly, even when it does not benefit expected gain (53).

In our sensorimotor confidence task we asked participants to report their confidence in their success at hitting the target, while performing a visuomotor adaptation task. Participants reached to a target straight ahead of them while experiencing error-clamped feedback, which they knew to be moving in a fixed direction that was not controlled by their own movements. Although the goal of the reach remained the same for all trials, every participant showed the effects of implicit adaptation, with the direction of their reach moving away from the clamped feedback. The error clamp (black line, Fig. 6) followed a 12 cycles per session sinusoidal time course, so the same frequency was expected for the reach direction (teal line, Fig. 6), which is the case for all participants.

**Fig 6:**
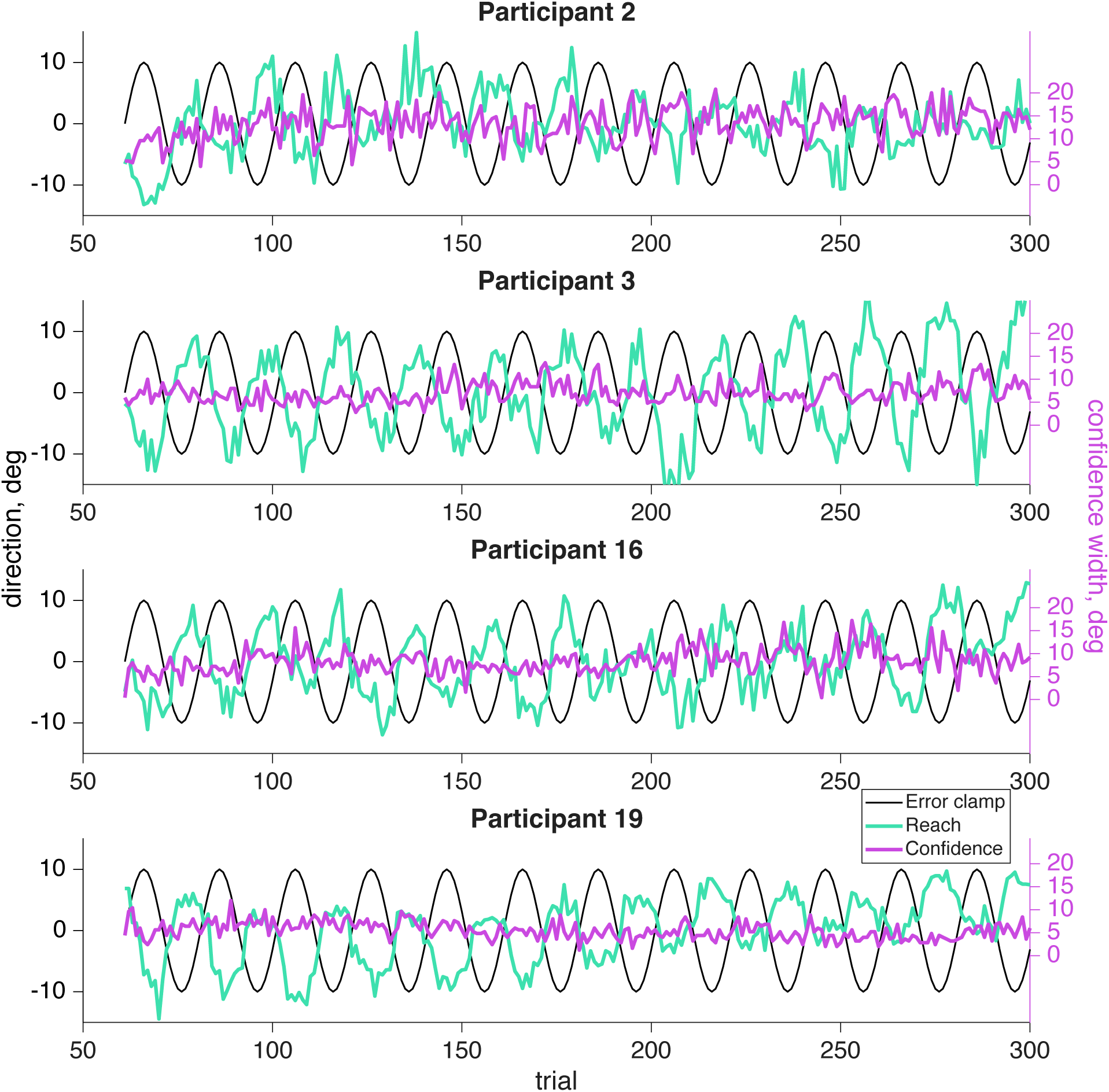
Sensorimotor confidence raw data. The error-clamp feedback (black line) was fixed by the experimenter and completed 12 cycles per session for the same sample participants as in Fig. 5. The participant was told to ignore the visual feedback and reach directly at the target, fixed at 0°. However, due to implicit adaptation the participant’s reach (teal line) followed the 12 cycles per session frequency of the error clamp, but was shifted by an average 11.89 trials. The confidence reports (magenta line) are always positive and have their own *y*-axis scale on the right side of the plot. The average confidence report is aligned with the left *y*-axis 0° point.

After every reach participants adjusted the width of a confidence arc (magenta line, Fig. 6) centered on the target location to attempt to enclose their true reach direction. Note, the confidence report will always be positive as it represents the magnitude of the arc. If they were very confident their reach landed near the target, then they should report a small arc. However if they felt they were far from the target, or unsure about their reach direction, it would be more advantageous for a larger arc to be used to maximize expected gain. We expected low confidence when either the error clamp or the reach itself was far from the target, which happens twice for each cycle of the error clamp (both clockwise and counterclockwise). As a result, we should expect to see a 24 cycles per session frequency in their confidence if it is affected by the error clamp or reach direction. If confidence was based purely on prospective information (e.g., knowledge of one’s motor uncertainty), we would not expect to find a significant 24 cycles per session frequency component.

As has been widely shown previously, adapted responses lag the perturbation (here, the error clamp) by several trials. Also, the adaptive response is to reach in the opposite direction from the perturbation in attempt to bring sensory feedback to its predicted position (Fig. 7A). Since there are 20 trials in one full cycle of the error-clamp direction, true anti-phase would result in a reach lag of 10 trials (i.e. on trial 11 the reach went in the same direction as the error clamp on trial 1). When the motor-learning response lag and anti-phase correction are both taken into account we expect the adaptive reach to lag to be just over half a cycle behind the error clamp, that is, more than 10 trials (37, 54–57). As expected, the reach direction lag seen in this sensorimotor task was always larger than 10 trials, on average 11.89 trials (SD: .61 trials) away from the error clamp (Fig. 8). This means motor-learning contributed a 1.89 trial lag past a true anti-phase lag with the error-clamp feedback.

**Fig 7:**
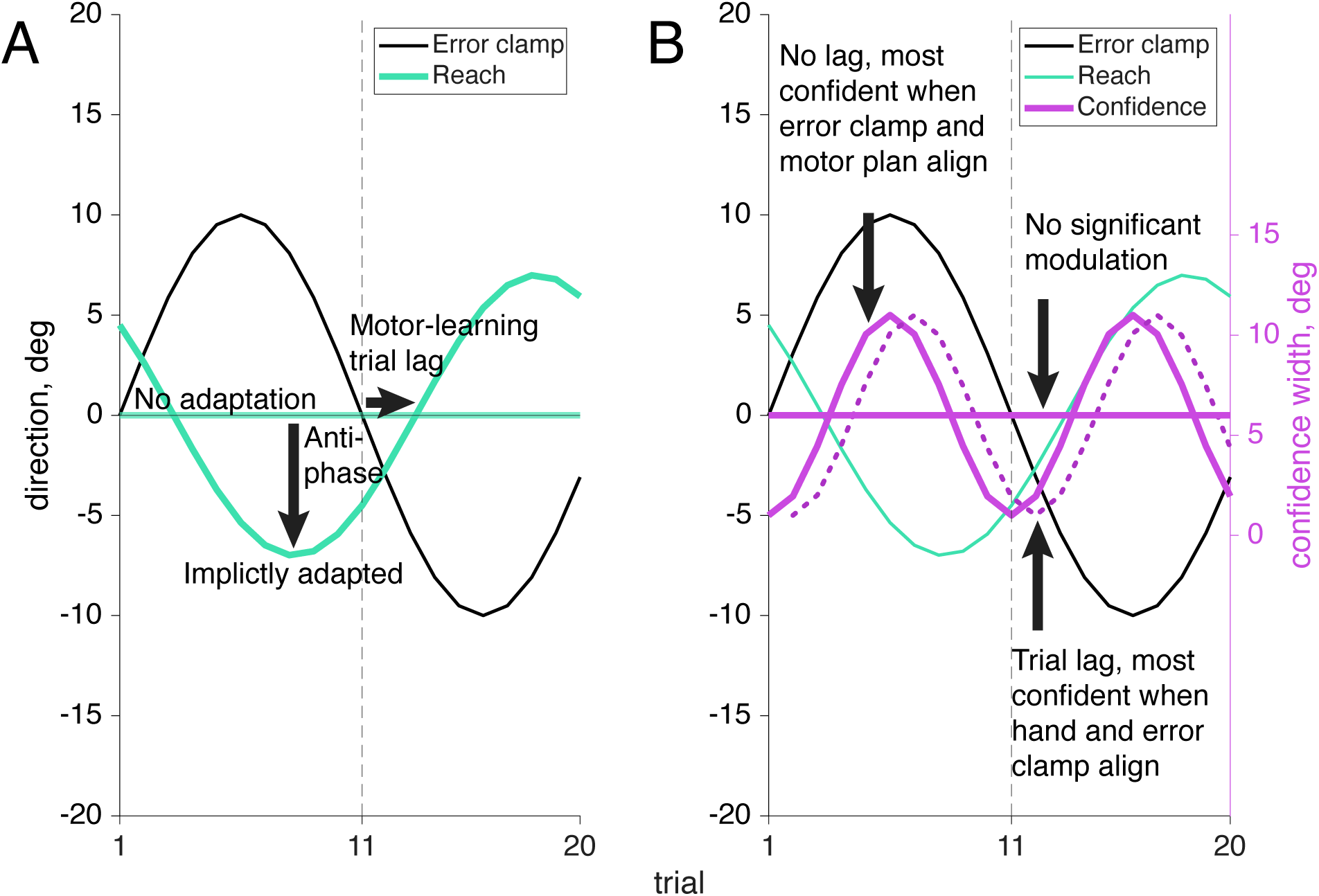
Example outcomes for reach and sensorimotor confidence response to the error clamp. A. When the error-clamped visual feedback is turned on (black line) displayed feedback changes its direction incrementally in a sinusoidal pattern across trials. If the participants fully ignore the feedback (which is known to be false) no adaptation would be shown and they would continue to reach for the target at 0°. Alternatively if implicit adaptation is present we will see the reach direction in the opposite direction from the error-clamp feedback (anti-phase) with a lag of a few trials to account for motor learning. B. Points are awarded when the true reach direction is enclosed in the confidence arc. No significant modulation would be present if the participant always assumed they were reaching straight forward at the target (solid line at 0°). If visual feedback was the primary influence on confidence, the width of the arc would fluctuate in sync with the error clamp (known to be false), leading to the highest confidence (smallest arcs) when the error clamp was close to the target and thus aligned with the motor plan, and lowest (largest arcs) when the distance from the target to the error clamp was greatest (solid cosine wave). Alternatively if proprioceptively sensed hand position contributed to the confidence rating as well, we’d expect to see the highest confidence (smallest arc) when the visual feedback and hand position are aligned (dotted cosine wave).

**Fig 8:**
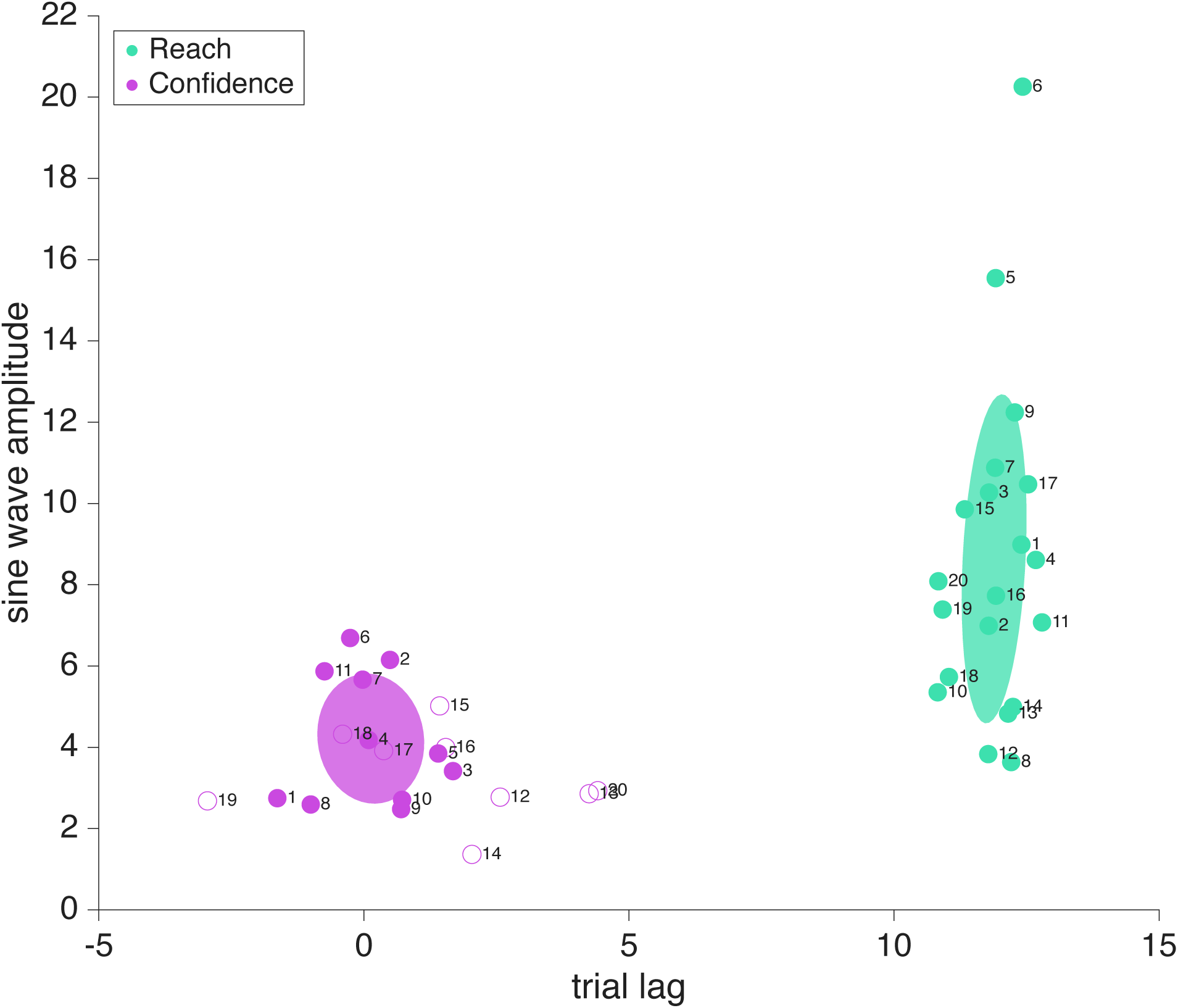
Sensorimotor confidence Fourier components for four sample participants. Trial lag and adaptation amplitude shown with closed dots represent participants that had a significant frequency component (24 cycles per session in the confidence reports and 12 cycles per session for the reaches), and open dots represent those who did not. Significance was determined using a shuffled permutation test of the Fourier analysis. All participants’ reaches showed a significant 12 cycle per session frequency component. The error-clamp direction (i.e., 0 lag) was determined by the experimenter and always followed a sinusoidal trajectory across trials with an amplitude of 10°. Reach direction (teal dots) lags relative to he error clamp were nearly constant across participants. However, about a third of the participants had adaptation amplitudes greater than the error-clamp amplitude of 10°. Confidence reports (magenta dots) follow a negative cosine trajectory and had an average trial lag of 0.13 when significant. The covariance elapses shown with the data represent only the participants with significant frequency components (filled dots).

Confidence judgments on each trial could be based on any combination of information available to the participant, from prospective information such as target location and knowledge of one’s own motor noise, to retrospective information including visual feedback and proprioception (Fig. 7B). The target location, which directs the motor plan, is fixed across trials and is available prior to the reach, as is an internal measurement of one’s own motor uncertainty. Retrospectively, additional information may be taken into account, such as visual feedback, which is known to be false but still induces implicit adaptation in all participants, and proprioception, the sense of where the hand is positioned in space, which varies from trial to trial. Previous research has shown that some individuals depend heavily on prior information while others incorporate all available data to maximize their performance (6). Participants 1 – 11 showed a significant (*p*<.01) 24 cycles per session frequency component in their confidence reports, based on the permutation test, while participants 12 – 20 did not show this significant component. The participant number order was arranged post-hoc to group participants by their phase significance in order to improve visual interpretation of the results, and was not the order the participants were originally tested in.

For participants whose confidence report demonstrated a significant frequency component of 24 cycles per session (participants 1-11, Fig. 8, magenta filled dots), there was a bit more variation in trial lag for peak confidence than in the trial lags for reach adaptation. When the target frequency of 24 cycles per session was significant, confidence lag was on average 0.13 trials (SD: 1 trial) beyond the error-clamp visual feedback. Because confidence can never be negative, rightward and leftward directions of the error clamp should have equivalent effects on confidence, resulting in a frequency doubling to 24 cycles per session. Additionally, as confidence is highest when the reported arc magnitude is small (the valleys of the wave), confidence follows a negative cosine structure. We assume a phase-zero baseline of maximum confidence when the error-clamp feedback is aligned with the target location. This also means that the wavelength is shorter for confidence than for the error clamp, meaning a lag of 5 trials would put it in anti-phase to the error clamp, as opposed to the 10 trial lag seen in the anti-phase reaches. Lags less than 0 trials mean that the peak of confidence arc size (i.e. lowest confidence) precedes the largest clamp distance, while lags greater than 0 trials show that confidence is lowest after viewing the greatest clamp distance. When there is a 10 trial difference between the confidence-arc lag and the reach-direction lag (confidence arcs get larger when the hand is further from the target) we can infer that proprioception is being used (e.g., if the average confidence lag was 2 trials while reach adaptation was 12). Alternatively, if there is a 0 trial difference between the confidence arc and the error clamp, confidence is highest (small arcs) when the feedback is near the target, regardless of hand position, e.g., if the trial lag was around 0, as we see in Fig. 8 (magenta filled dots). We can infer confidence is being affected by the the same system that is separately influencing implicit adaptation, even when the clamped feedback is known to be false.

For participants who did not show a significant 24 cycles per session frequency in confidence (participants 12-20, Fig. 8 open dots), the estimated lags are random and it is likely they are primarily focusing on the motor goal of the target location and prior knowledge of their own motor noise, actively ignoring trial-specific information such as the false visual feedback and proprioceptive signals. We’ve included the amplitude and trial lag at 24 cycles per session from the Fourier analysis as these values are recoverable, although the frequency was not statistically significant using the permutation test. They are denoted with open dots in Fig. 8. For the participants who did not show the significant 24 cycles per session frequency in confidence, the mean confidence rating still remains consistent across trials so the mean line has been plotted (Fig. 9).

**Fig 9:**
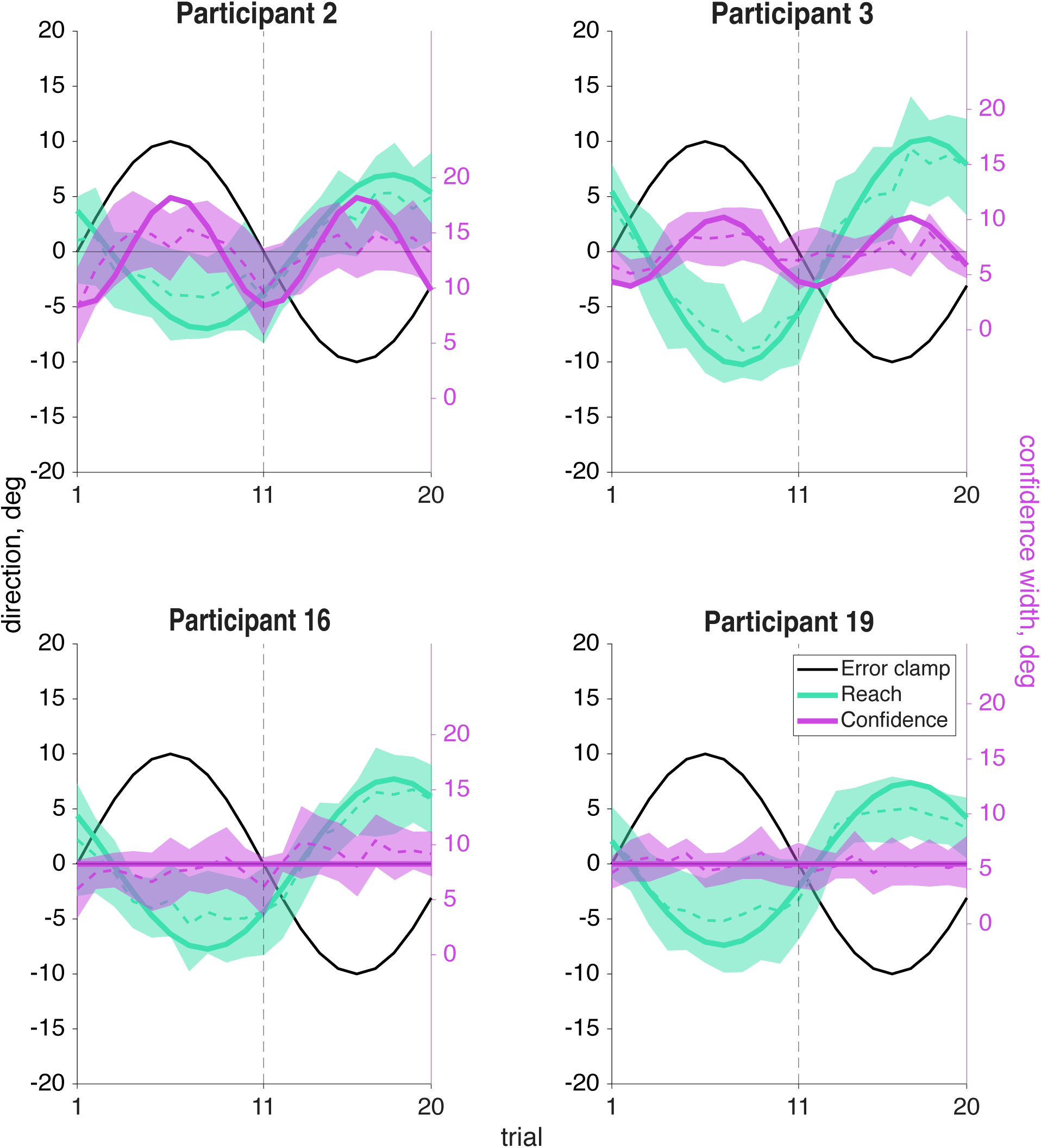
Sensorimotor confidence sine-wave fits. An averaged cycle (dashed line with shaded CI) and an overlayed sine wave (solid line) using the phase and amplitude components from a Fourier analysis at the expected frequency of the response is shown here for the same four sample participants. Participants 2 and 3 on the top row had a significant 24 cycles per session frequency in confidence (magenta), while participants 16 and 19 on the bottom row did not show this significant frequency component. The sine wave has phase and amplitude from a Fourier analysis at the frequency of 24 cycles per session when that amplitude was significantly different from zero (Participants 2 and 3). For the other participants the mean is plotted (i.e., 0 amplitude). All participants showed a significant 12 cycles per session frequency for reach direction (teal). Plots for all participants are in Supplement 2.

When the 24 cycles per session frequency in confidence was significant, the trial lag was such that confidence was not always lowest when either the error-clamp feedback or the hand-position error was at its maximum, suggesting that confidence was in response to some combination of the two cues. If both visual feedback and hand position were taken into account, it’s possible that the mismatch between the two sources of information led to a reduction in confidence. We observe a significant correlation (significant *r values*, *p*<.05, *N*=10, Fig. 10A) between confidence reports and the distance between the feedback direction and the hand direction in half of our participants (participants 1-8, 14, 18, Fig. 10B), a subset that is highly overlapping with the participants displaying a significant 24 cycles per session frequency component in confidence.

**Fig 10:**
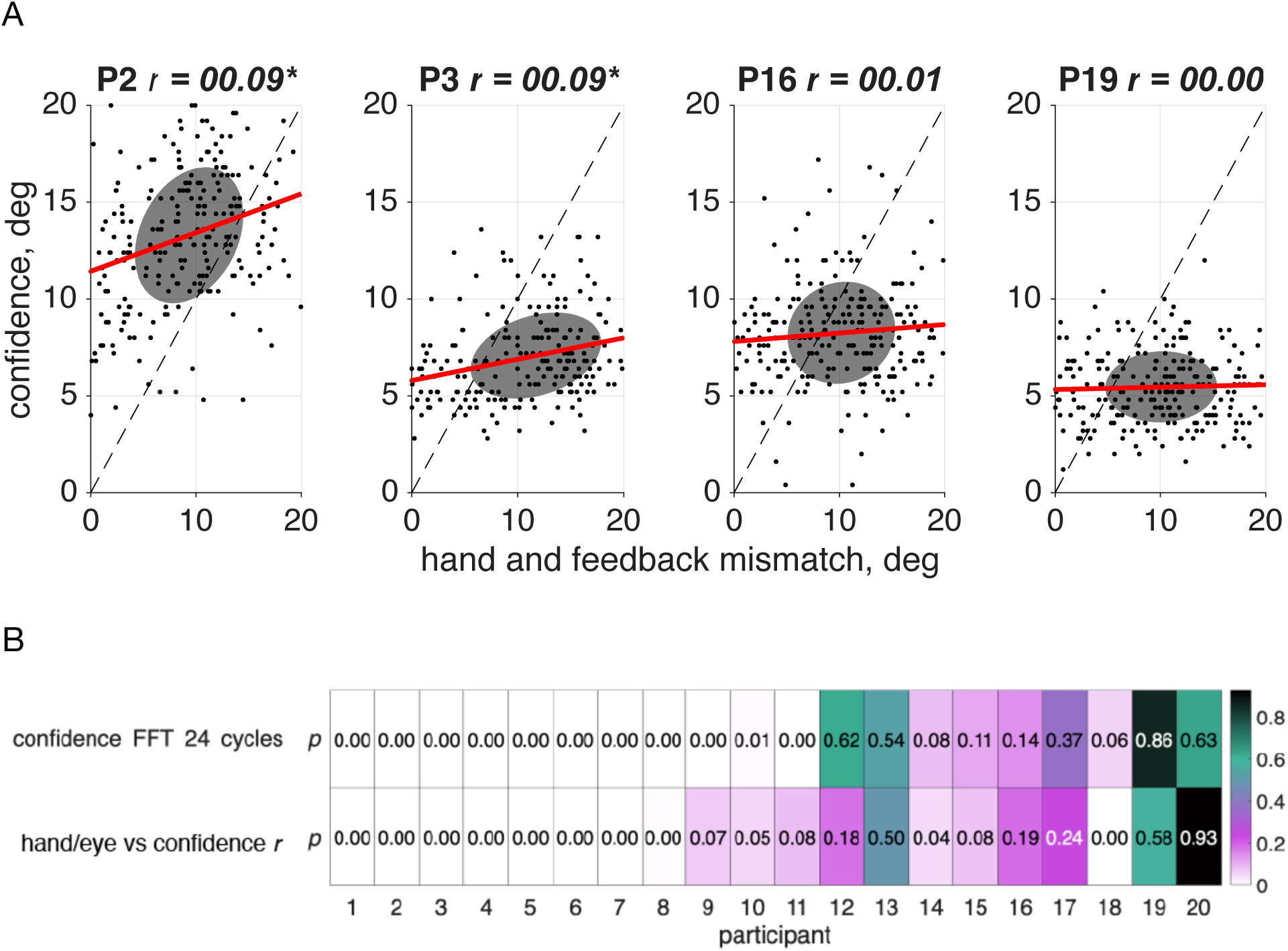
Correlation between sensorimotor confidence and the mismatch between hand and feedback direction. A. Correlation between the confidence report on a given trial and the distance (in degrees) between the visual feedback location and hand position for the same four sample participants. Since the reach lag is on average 1.89 trials past true anti-phase, the point at which the mismatch is the smallest occurs when the error clamp is slightly off center from the target. Plots for all participants are in Supplement 3. B. Participants who showed a significant 24 cycles per session frequency component in their confidence judgments were more likely to demonstrate a significant correlation, meaning that the greater the mismatch between the visual error-clamp feedback and their sensed hand position, the less confident they would be that they hit the target.

On each trial participants had the opportunity to earn points with their confidence judgment. Participants in this task earned an average of 540.38 points (SD: 265.36) during the session, and successfully captured their true reach direction on an average of 39.3% of trials (SD: 18.6%). Comparing the average points earned between participants that showed a significant 24 cycles per session frequency in their confidence reports and those who did not, there was no significant difference between the two groups, (*t*(18) = 1.54, ns). This indicates that while strategies may differ as to their use of proprioception and prior experience, there was not a large benefit points-wise from modulating the confidence reports (i.e., showing a significant frequency component of 24 cycles per session).

### Motor-Awareness Confidence

In our motor-awareness confidence task we asked participants to report perceived reach direction and their confidence in how accurate that estimate was, instead of how successful they were at reaching the target. This report direction (tan line, Fig. 11) is similar to the confidence judgment in the sensorimotor-confidence task in that it is a judgment made about the reach that was just performed. However, while the sensorimotor-confidence judgment used an arc that was centered on the target and extended uniformly in both directions, the motor-awareness report was a single point that could be rotated in any direction. Because this motor-awareness judgment explicitly asked about the position of the hand, the motor plan or proprioception must be used, although it is not fully necessary to incorporate it into the sensorimotor-confidence judgment. After the sensed direction was reported, an arc was adjusted to report confidence centered on the reported direction. This judgment specifically asks the participant to report how confident they were that their reported direction was their true reach direction. To maximize point gain in the sensorimotor-confidence task the confidence-arc size needed to expand with the adaptation magnitude (points were only earned on trials where the true reach direction was captured), however for the motor-awareness confidence judgment the magnitude of the arc does not need to fluctuate with the adaptation to maximize points so long as the report direction was accurate.

**Fig 11:**
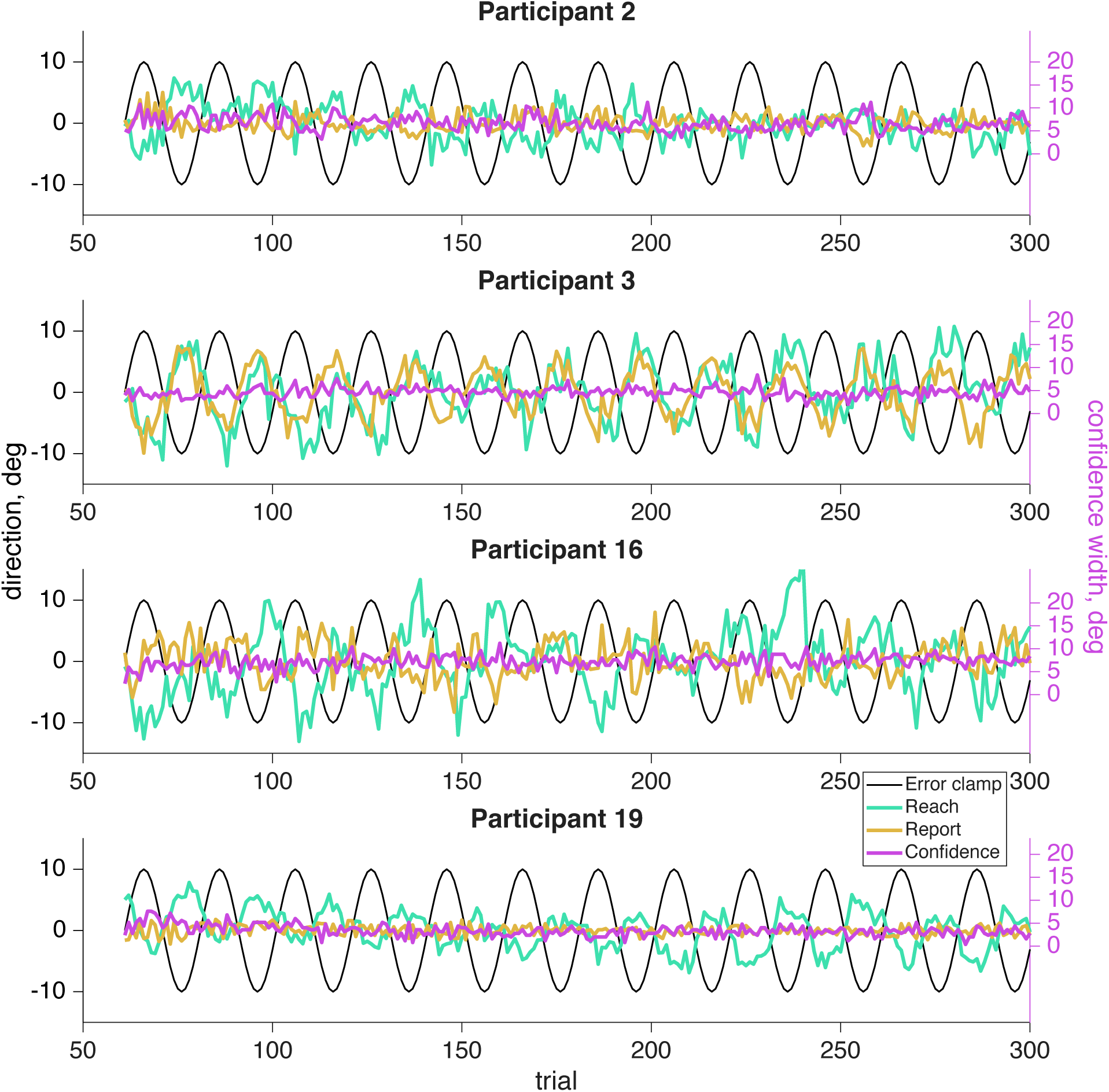
Motor-awareness confidence raw data for four sample participants. The error-clamp feedback (black line) was fixed by the experimenter and completed twelve cycles per session. The participant was told to ignore the visual feedback and reach directly at the target, fixed at 0°. However, due to implicit adaptation the participant’s reach (teal line) followed the 12 cycles per session frequency of the error clamp, but lagged the error clamp by an average of 11.84 trials. The reported sensed hand position (tan line) also may contain the 12 cycle frequency. The confidence report in the success of the reported hand position (magenta line) could never be negative so it has its own *y*-axis scale on the right side of the plot. The average confidence report on the right-hand scale is aligned with the left *y*-axis 0° point.

If the report direction was influenced by the implicit adaptation, a significant frequency component of 12 cycles per session would be present in the reports (Fig. 12A). Out of the 20 participants 16 showed a significant (*p*<.01) frequency component at 12 cycles per session (participants 1-7, 9, 11-18; Fig. 13, tan filled dots). Participants who showed a significant (*p*<.01) 24 cycles per session frequency in confidence (participants 1-9, 12-13, Fig. 13, magenta filled dots) indicate that confidence in one’s hand position was influenced by conflicting signals; the motor goal, visual feedback, and proprioception (Fig. 12B). This is similar to what was seen in the sensorimotor-confidence task suggesting that similar sensory inputs influence confidence in the success of a sensory directed goal as well as motor awareness.

**Fig 12:**
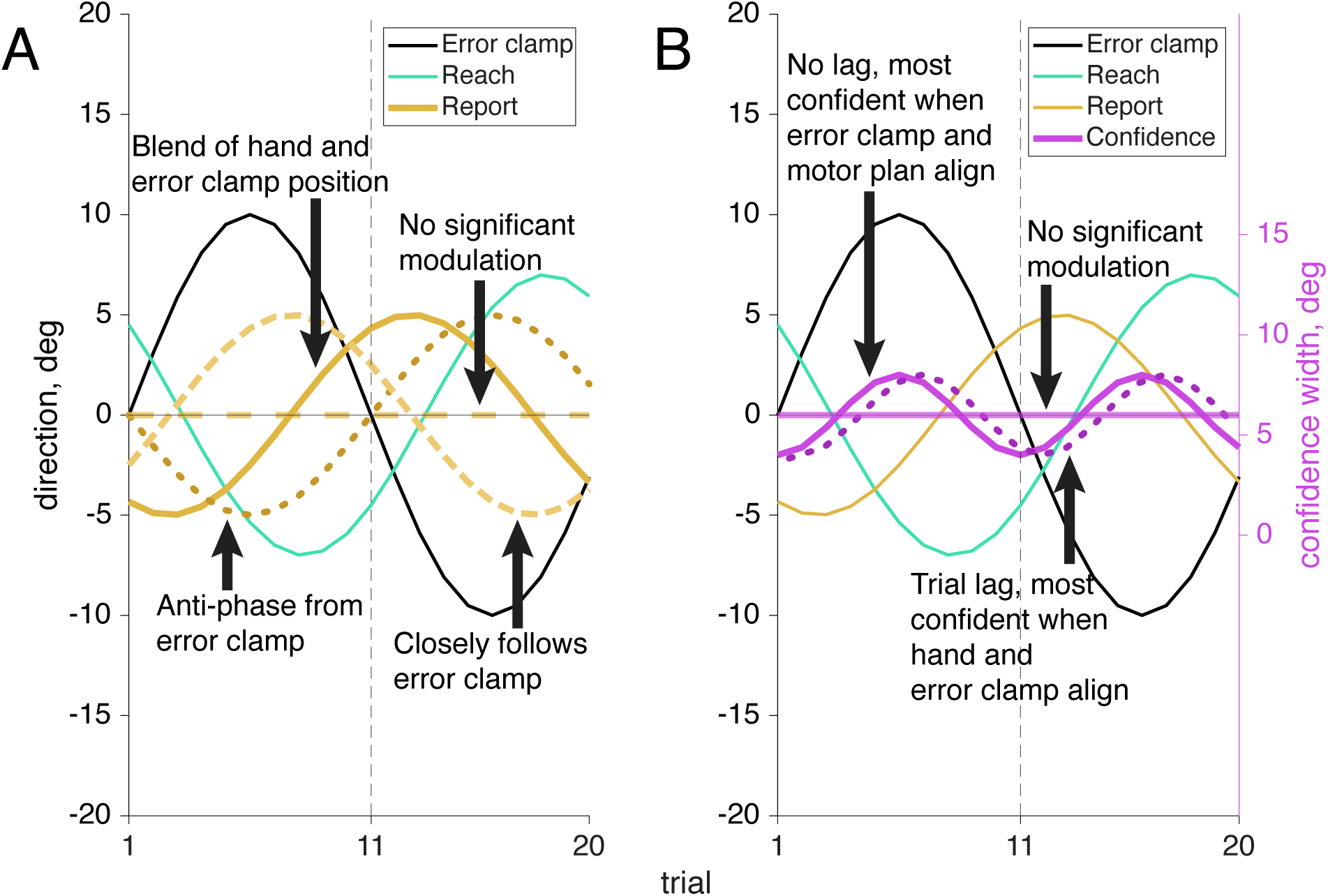
Example outcomes for report and motor-awareness confidence response to error-clamp driven implicit adaptation. A. If there is no conscious influence of the error-clamp or proprioceptively sensed hand position on the report (known to be false or uncertain respectively) then the participant may follow their motor plan, and report their endpoint at the target location (dashed line at 0°). Alternatively the participant could be greatly influenced by the error clamp and report their reach in the same direction (dashed sine wave). Although if they are aware of some adaptation happening, but their proprioception is known to be uncertain, they may report their reach direction anti-phase to the error clamp, but with a smaller magnitude as the motor plan still has influence (dotted sine wave). Lastly they could be combining the two signals and end up with a report somewhere in the middle (solid sine wave). B. Since the arc is centered on the reported location and the ability to score points is based on whether the reach direction was enclosed in the confidence arc modulating the confidence judgment is not necessary to perform well in this task. The success of the reported direction is not contingent on its distance from the target so we would expect no significant modulation here (solid line at 0°). However, if the error clamp heavily influenced the confidence judgment we would expect to see highest confidence (smallest arcs) when the error clamp is close to the target (solid cosine wave). If the hand position is also taken into account we would expect to see the highest confidence when there is minimal mismatch between the error-clamp feedback and the hand position (dotted cosine wave).

**Fig 13:**
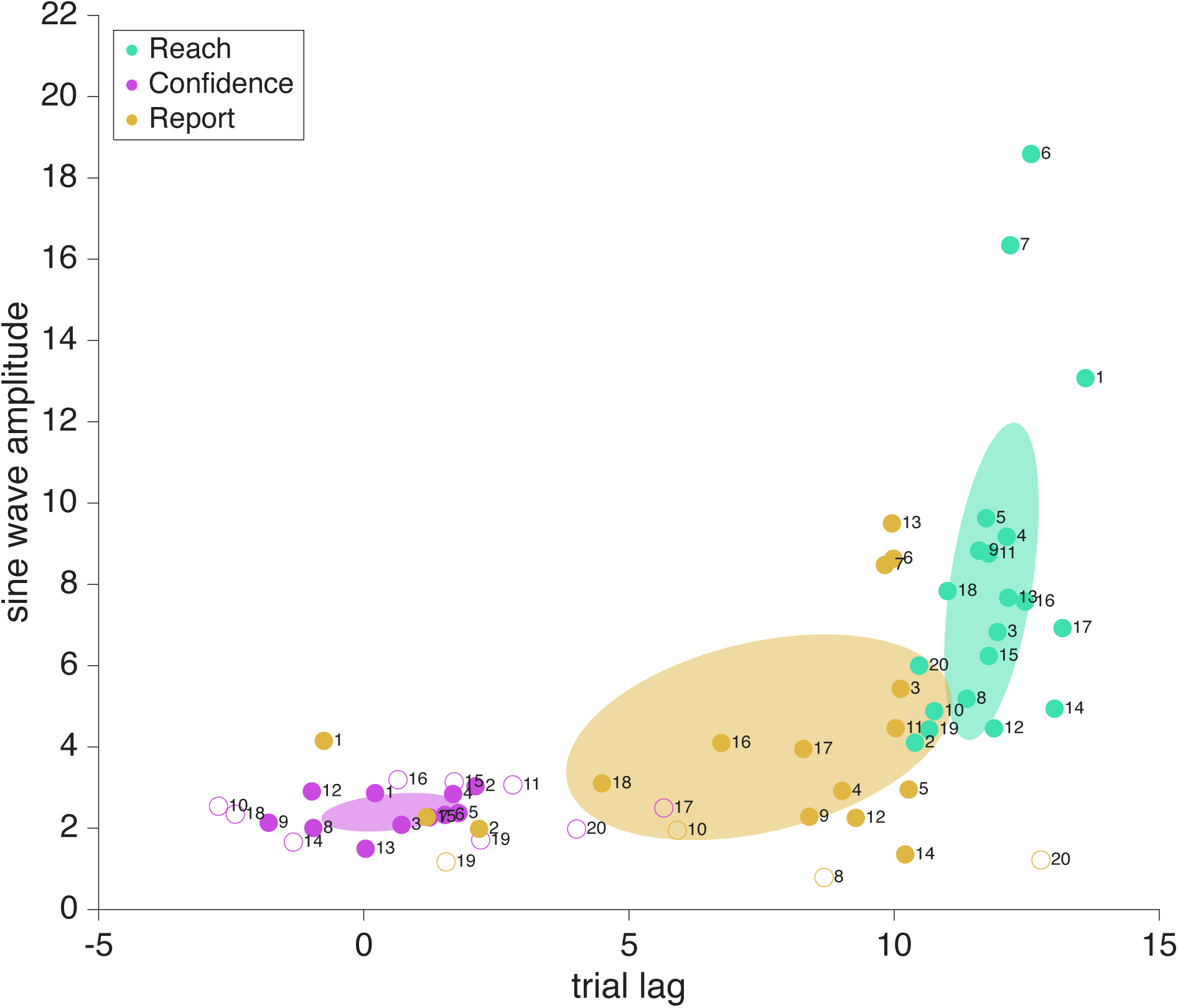
Motor-awareness confidence Fourier components. Lag and adaptation amplitude shown with filled dots representing participants that had a significant frequency component (24 cycles per session for confidence, and 12 cycles per session for reaches and reports), and open dots for those who did not. Significance was determined using a shuffled permutation test. All participants’ reaches showed a significant 12 cycle per session frequency component. The error-clamp direction (i.e., 0 lag) was determined by the experimenter and always followed a sinusoidal trajectory across trials with an amplitude of 10°. Reach-direction lags (teal dots) were nearly constant across participants, however a few showed adaptation amplitudes greater than the error-clamp amplitude of 10°. Confidence reports (magenta dots) had an average trial lag of 0.51 when significant, while the adaptation lag in the reported hand position (tan dots) was much more variable across participants. The covariance ellipses shown with the data represent only the participants with significant frequency components (filled dots).

While the trial lags for reach direction and significant confidence reliably clustered around 11.84 (SD: .89 trials) and 0.51 (SD: 1.3 trials) respectively, the trial lag for report direction was far more variable (M: 7.45, SD: 3.64 trials) when a significant frequency of 12 cycles per session was present (Fig. 13, tan filled dots). Notably, most of the participants had a report lag less than than 10 trials, and all participants had a reach adaptation trial lag of over 10. This indicates a strong influence of the visual feedback on reported hand position as it was uncommon for participants to estimate the hand position with a lag beyond true anti-phase.

The adaptation lag and amplitude of the report varied across participants (Fig. 13, tan dots), with four major patterns presenting themselves in the data (Fig. 14). Participants with a report lag from 8 – 10 trials (participants 3-9, 12-14) were anti-phase to the error-clamped visual feedback and followed the leftward / rightward direction of the hand position well but did not capture the delay. In opposition to that, participants 1, 2 & 15 showed a report adaptation lag centered around 0 trials with their report direction pulled toward the visual feedback and away from the true hand position. In between the two we had participants 16-18 with a trial lag from 4 to 8 where the reported position seemed to be influenced by both the feedback and hand directions. Lastly we had four participants who didn’t show a significant 12 cycle per session frequency component in their reports (participants 8, 10, 19 & 20) and kept their reported position near the motor-goal direction, ignoring both the feedback and their sensed hand position.

**Fig 14:**
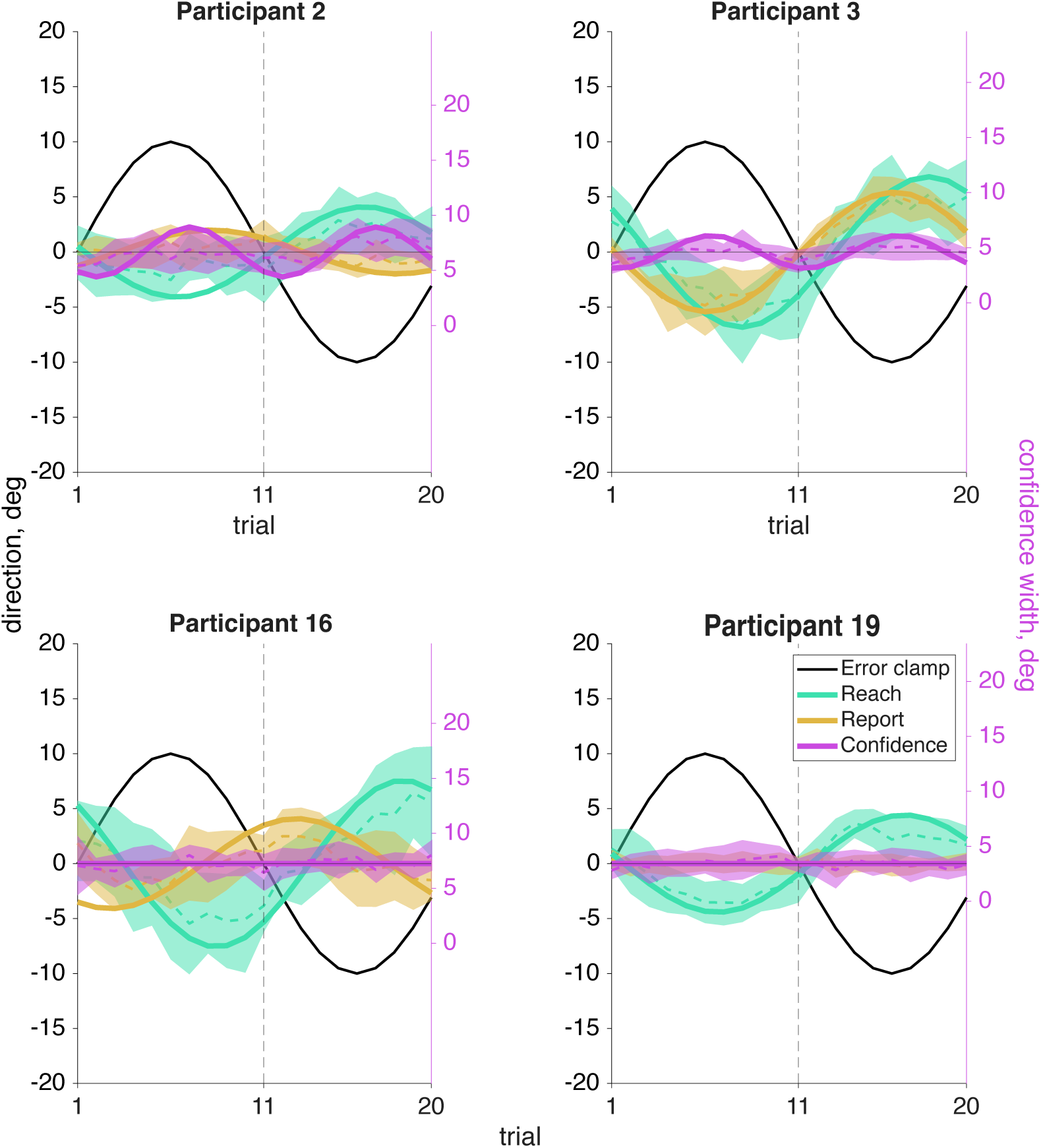
Motor-awareness confidence sine wave fits. An averaged cycle (dashed line with shaded CI) and an overlayed sine wave (solid line) using the phase and amplitude components from a Fourier analysis at the expected frequency of the response is shown here for the same four sample participants. Participants 2 and 3 on the top row had a significant 24 cycles per session frequency in confidence (magenta), while participants 16 and 19 on the bottom row did not show this significant frequency component (and so the sine is plotted at zero amplitude). All participants showed a significant 12 cycles per session frequency for reach direction (teal). Participant 19 is the only one to not show a significant 12 cycles per session frequency in the reported hand position (tan), but the other three sample participants here all show notably different trial lags of the report relative to the error clamp. Plots for all participants are in Supplement 4.

Confidence-arc amplitude was smaller in the motor-awareness task than in the sensorimotor task. This is because once the reach direction was indicated the participant had already accounted for the distance from the target. The motor-awareness confidence judgment only represented their uncertainty in that report. This reduced much of the variance seen in the sensorimotor confidence reports, which had a fixed mean at the target location so larger reports were needed when the hand position was sensed far away. In the sensorimotor confidence task a large arc would be required if the participant felt they had missed the target by a great distance. However in the motor-awareness confidence task the arc could be centered on where they sensed the hand to be, even if it was far from the target. The confidence arcs reported in the task were smaller overall because of this.

As in the sensorimotor-confidence task we found that just over half of our participants (Fig. 15B) demonstrated a significant correlation (significant *r* values, *p*<.05, *N*=11) between confidence and the mismatch between their hand and feedback directions. Notably only five participants (1, 2, 6, 7, & 10) had a significant correlation (significant *r* values, *p*<.05, *N*=5, Fig. 15A) between confidence and the distance between their report and true reach direction, which is the metric that the points are based on. While only proprioception is needed to indicate perceived reach direction, in our task the motor goal and the error-clamped visual feedback both influenced the reported direction and the confidence in its success.

**Fig 15:**
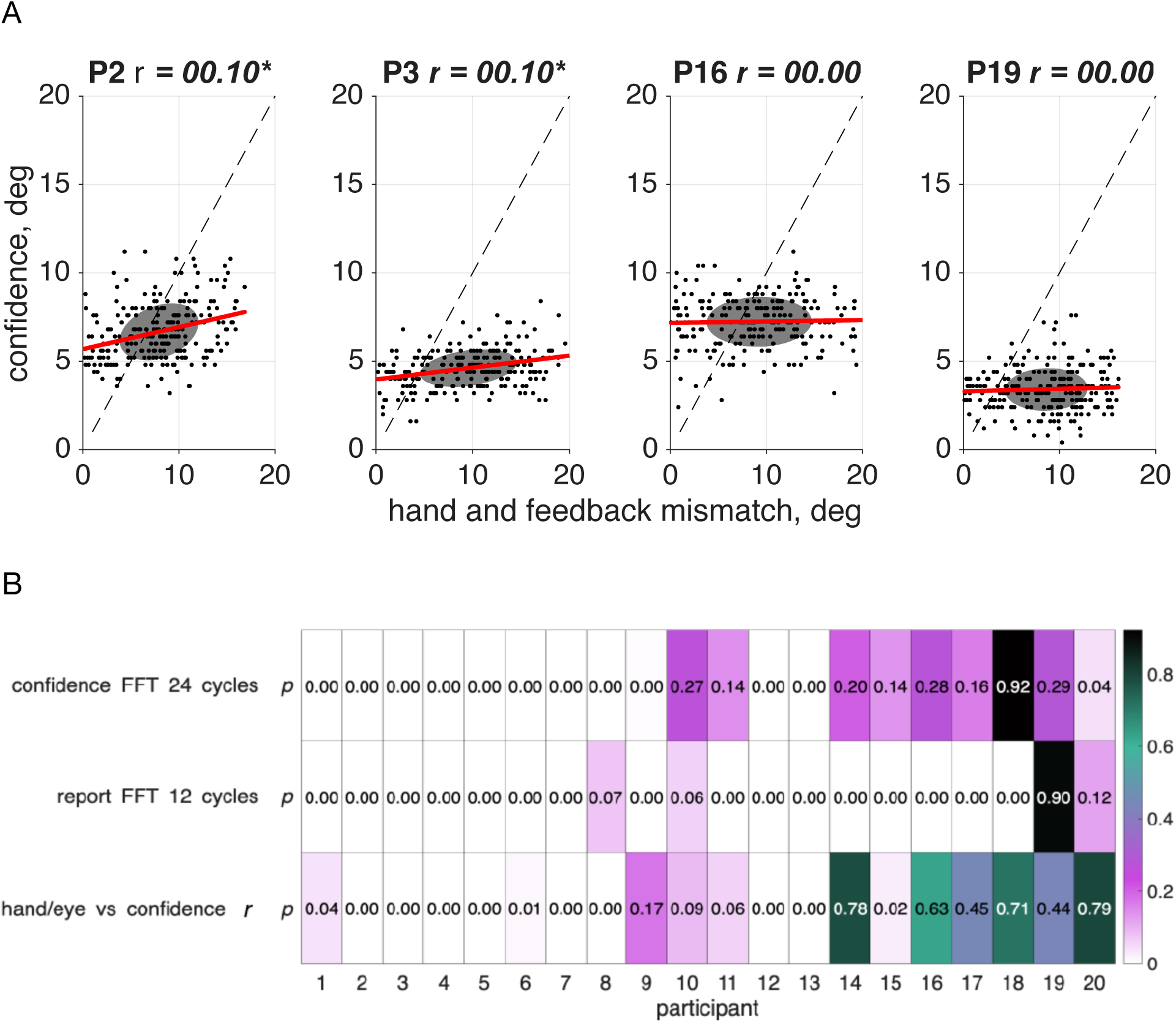
Correlation between motor-awareness confidence and the mismatch between hand and feedback position. A. Correlation between the confidence report on a given trial and the distance (in degrees) between the visual feedback location and hand position for the same four sample participants. Since the reach lag is on average 1.84 trials past true anti-phase, the point at which the mismatch is the smallest is not centered at the target position, 0°. Plots for all participants are in Supplement 5. B. Participants who showed a significant 24 cycles per session frequency component in their confidence judgments were more likely to demonstrate a significant correlation, meaning that the greater the mismatch between the visual error-clamp feedback and their sensed hand position, the less confident they would be that they hit the target. In this experiment the confidence report is in the success of the reported hand position, and shouldn’t be influenced by the distance from the error clamp, however in over half the participants we see that there is a significant correlation between the two.

In this task as well, participants had the opportunity on each trial to earn points with their confidence judgment. Participants in this task earned an average of 294.44 points (SD: 155.91) during the session, and successfully captured their true reach direction on an average of 27.0% of trials (SD: 15.4%). Overall performance was significantly worse in this task than in the sensorimotor-confidence task, *t*(19) = 4.64, p<.000, when it came to maximizing the expected point gain, with participants earning an average of 245.94 (SD: 237.25) fewer points. Comparing the average points earned between participants who showed a significant 24 cycles per session frequency in their confidence reports and those who did not, there was a significant difference between the two groups, *t*(18) = 3.46, p<.01), with participants who showed the significant frequency component (participants 1-9, 12 & 13, M: 207.64, SD: 137.00) scoring significantly lower than those who did not show that significant frequency component (participants 11, 10, & 14-20, M: 400.53, SD: 106.07). There was no significant difference between these groups in their average proximity difference between the reported reach direction and the true reach direction, so this difference in points is driven by the confidence judgments, not by the report accuracy.

### Cross-Experiment Comparisons

We compared the proprioceptive uncertainty measured in the separate motor-awareness task to confidence and report trial lags as well as the adaptation amplitude (Fig. 16). There was no significant difference between the magnitude of the proprioceptive uncertainty for participants who had a significant 24 cycle per session frequency component for confidence judgments and those who did not. This was true both for sensorimotor confidence judgments (*t*(18) = .56, *ns*) and motor-awareness confidence (*t*(18) = 1.4, *ns*).

**Fig 16:**
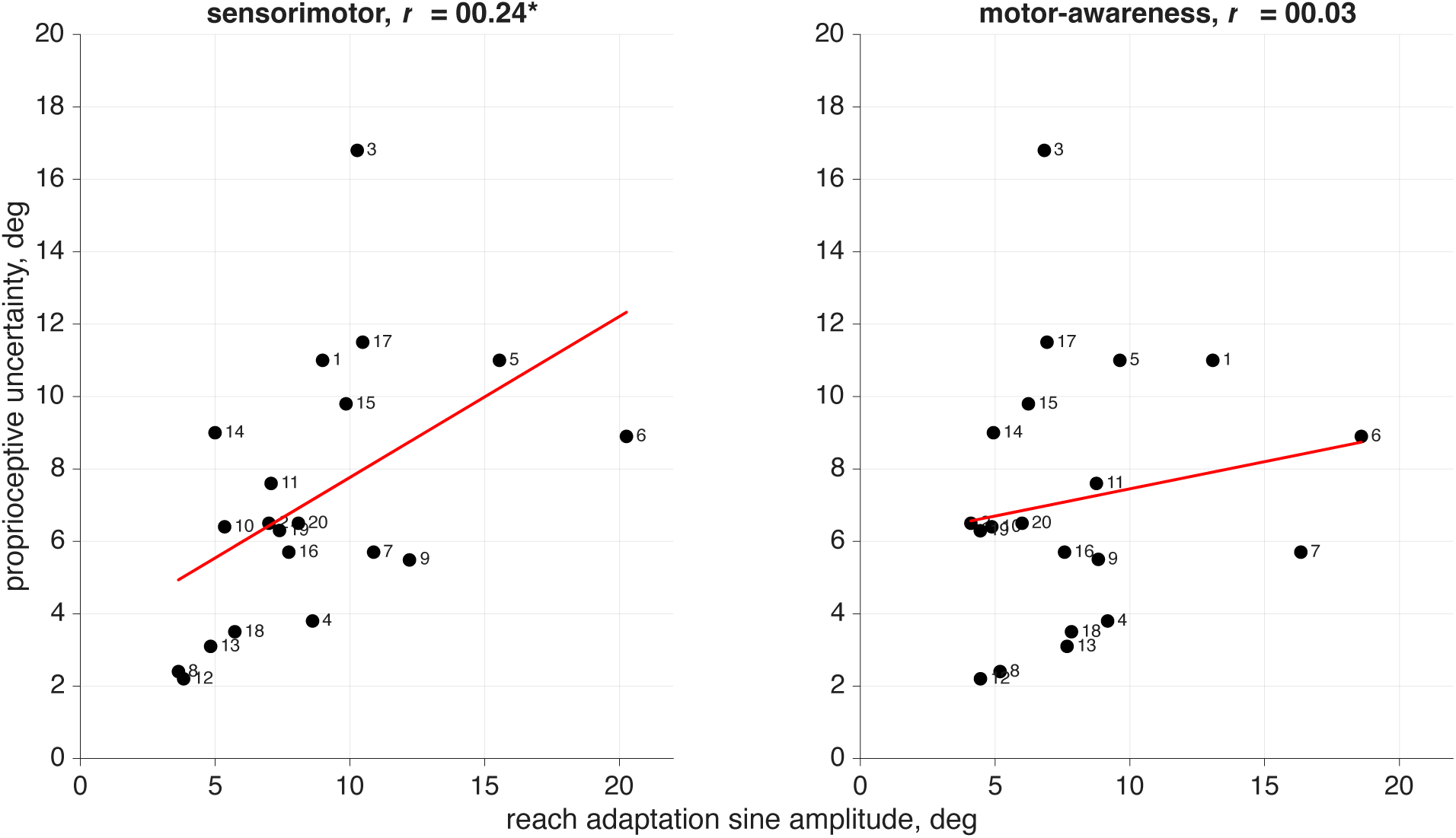
Motor-awareness proprioceptive uncertainty compared with reach-adaptation sine wave amplitude. Correlation comparison between the proprioceptive uncertainty value measured in the separate motor-awareness task, and the reach-adaptation amplitude at the 12 cycles per session Fourier component from the sensorimotor confidence and motor-awareness confidence tasks. There is a significant correlation between proprioceptive uncertainty and reach amplitude during implicit adaptation when making sensorimotor-confidence judgments but not when performing motor-awareness reports.

There was a significant correlation between reach-adaptation amplitude and proprioceptive uncertainty in the sensorimotor-confidence task (*r* = .24, *p*<.05). However, this significant relationship was not present in the motor-awareness confidence task (*r* = .03, ns). We assume that participants with a better sense of proprioception (lower uncertainty) would adapt less than those with higher uncertainty, which is what we are seeing in the sensorimotor task. Those who are more aware of their hand position would be more likely to sense the implicit adaptation and attempt to consciously correct for it to hit the target. It is possible that the inclusion of the report in the motor-awareness task made participants focus on their hand position more intently, reducing the amount of adaptation that was seen in the sensorimotor task when hand position wasn’t directly queried. There is a significant correlation between the reach adaptation amplitudes from the sensorimotor task and the motor-awareness task within participant (*r* =.52, p<.000), indicating that participants behaved similarly in both tasks.

## DISCUSSION

In this paper we examined how both sensorimotor confidence and motor-awareness confidence are affected by implicit motor adaptation. Because implicit adaptation automatically drives changes in reaching direction without participants’ awareness, it is surprising that prior studies have not asked about the subjective experience, whether in terms of motor awareness or sensorimotor confidence, at least systematically.

To address this gap, twenty participants performed three reaching tasks: a motor-awareness task to independently fit proprioceptive uncertainty, and two confidence tasks that asked participants to report sensorimotor confidence and motor-awareness confidence during a visuomotor adaptation task with an error-clamped sinusoidal perturbation. The imposed sinusoidal perturbation pressured (engaged) participants to make trial-by-trial adjustments in their confidence reports and made possible frequency-based Fourier analyses to detect subtle changes and relationships between our variables of interest. Via permutation tests, we found significant frequency components for each participant’s reach direction (implicit adaptation), sensed hand position, and confidence reports, independently for the motor-awareness confidence and sensorimotor confidence tasks. This perturbation format uniquely allowed us to see adaptation on a trial-by-trial level throughout the whole session, and gave us a sensitive and objective way to relate changes in all measures, together forming a strong and innovative approach to a motor adaptation task.

### Individual differences in awareness

While all participants displayed a significant Fourier frequency component for reach direction, only half of participants showed significant components in their confidence judgments. This indicates that while the implicit adaption was successful for all participants, not all of them incorporated the shifted proprioceptively sensed hand position or error-clamped visual feedback into the confidence judgment.

While participants displayed robust adaptation, which followed similar trial-by-trial changes (with varying amplitudes), there were notable individual differences in their confidence reports. For the sensorimotor task about half the participants modulated their confidence reports in concert with the clamped visual feedback and/or sensed hand position, displaying a significant frequency of 24 cycles per session. A key question becomes why half of the participants modulated their confidence, while the other half did not.

One possibility is that participants varied in the degree to which they allowed sensory inputs to inform their explicit judgments. Because participants were explicitly told that the visual feedback was task-irrelevant and unaffected by their movement, some may have adopted a top-down strategy that treated both the clamped cursor and noisy proprioception as uninformative, so it’s possible they made a conscious effort to disregard visual and proprioceptive signals. The idea that participants might ignore proprioception is especially important for interpreting the sensorimotor task. In this task, the confidence arc was centered at the target location and there was an incentive to reach for the target. Participants who underestimated (or were unaware of) the effects of implicit adaptation may let their motor plan dictate a larger portion of their confidence judgment. With no veridical visual feedback of the true end point, it is possible that some participants only used information they knew to be valid. That is, they knew that the visual feedback was unrelated to their performance and the objective was to reach to the target. Thus, their motor goal was to aim toward the target, and therefore they may have been motivated to ignore any uncertain visual or proprioceptive information. Indeed, this can be seen when looking at the points metric for the task: participants only successfully captured their end point on 39.3% of trials, much lower than what was observed in previous research using a similar behavioral design (6, 34). Due to the implicit adaptation we expect participants to perform poorly on the points-based objective because they should not be aware of adaptation; if they were aware of adaptation (e.g., hand position drifting from the target), the objective to induce an implicit adaptation would have failed.

We see the greatest differences between participants in the sensed hand-direction reports from the motor-awareness confidence task. While the confidence judgments in the sensorimotor confidence task acted as a wager around how confident one was in having accurately hit the target, the report in the motor-awareness confidence task required the participant to specify a specific direction. The magnitude of the reported direction away from the target tended to be smaller than the true reach direction, potentially because participants assumed it was more likely for them to be closer to the target as that was their aiming direction, so centering the mean of the confidence judgment a little closer would have a higher probability of capturing the true end point.

Another difference between participants could be due to variability in susceptibility to proprioceptive adaptation (58–60). Some participants may be more prone to sense that their hand position is at the target, as this is their aiming location, recalibrating from the true adapted angle. This would mean that while a participant’s proprioceptive uncertainty may be low, they adapt where they sense their hand to be to counteract the implicit shift. In most real-world examples of implicit adaption, such as adapting to a new pair of eyeglasses, the small changes may not be noticeable to the observer. The reach goal should remain the same and the implicit adaptation should subconsciously shift the motor plan while the sense of reaching the target remains the same. It’s possible that this is the reason we don’t see strong relationships between the separately measured proprioceptive uncertainty and our confidence judgments in this implicit adaptation task, while these were indicated when no adaptation was present (6) and during explicit adaptation (34).

### Study Limitations

The error-clamp paradigm is successful at eliciting implicit adaptation, however it is not directly comparable to real-world sensory experience. It’s very unusual to have the visual feedback from a reach have its speed be controlled by your hand but the movement angle completely untethered from your movements. While the adaptation is automatic and stable across all sessions of all participants, the cognitive reports differ greatly between observers. The knowledge that the visual feedback is false and should be ignored may be taken to heart by some participants more than others, prompting them to make decisions based on top-down cognitive judgments about what they believe the best response should be, instead of relying on bottom-up sensory inputs that conflict with the motor plan. Because of the unnatural sensory mismatch found with the error-clamp paradigm, and the possibility of different cognitive strategies to gain points, the broad range of participant outcomes in the sensed hand-direction reports and confidence judgments could be expected.

The behavioral aspects of both tasks were nearly identical: a reach was made to a target positioned in front of the participant while they were shown error-clamped feedback, then they used the dial in their other hand to report confidence. However, they differed in what kind of confidence report was reported. In the sensorimotor confidence task we asked participants to report how confident they were that they hit the target, while in the motor-awareness confidence task we asked how confident they were in their self-reported perceived reach direction. Because of these different, reflections we expected the differences in magnitude present between the sensorimotor confidence judgments and the motor-awareness confidence judgments. In the sensorimotor confidence task the confidence report is a second-order judgment, the metacognitive decision about the outcome of an action. In the motor-awareness task however, the reported direction is the second-order judgment, and the confidence report becomes a third-order judgment ⏤ a metacognitive decision about the accuracy of an action’s outcome. Given that the confidence judgment is not directly about the hand position in this case, the presence of the 24 cycles per session frequency in some participants was unexpected. Notably, participants who showed this significant frequency scored significantly fewer points than those who did not, indicating that the information prompting this sinusoidal pattern in confidence ⏤ such as the error-clamp angle or reach-direction distance from the target ⏤ acted as a distraction from the important information here, which is the distance from the report to the reach direction. Participants who ignored this additional information scored more points on average.

### Conclusion

We used a reaching task to investigate how both sensorimotor confidence and motor awareness confidence are affected by an implicit adaptation. Confidence was reduced when there was a mismatch between the visual feedback and reach direction. This was expected with sensorimotor confidence as the performance metric is based on the target. However the influence of the adaptation still appeared in the motor-awareness confidence judgment although this judgment was not dependent on the distance from the target.

## Author Contributions

MHE carried out the formal analysis, investigation, software, data curation and visualization, and wrote the original manuscript. MHE, JAT and MSL collaboratively worked on conceptualization, methodology and editing. MSL and MHE are responsible for funding acquisition.

## Data Availability

All data and code files will be made available on OSF database and are hosted on GitHub https://github.com/marissahevans/implicitAdaptation

---

**Supplement 1:**
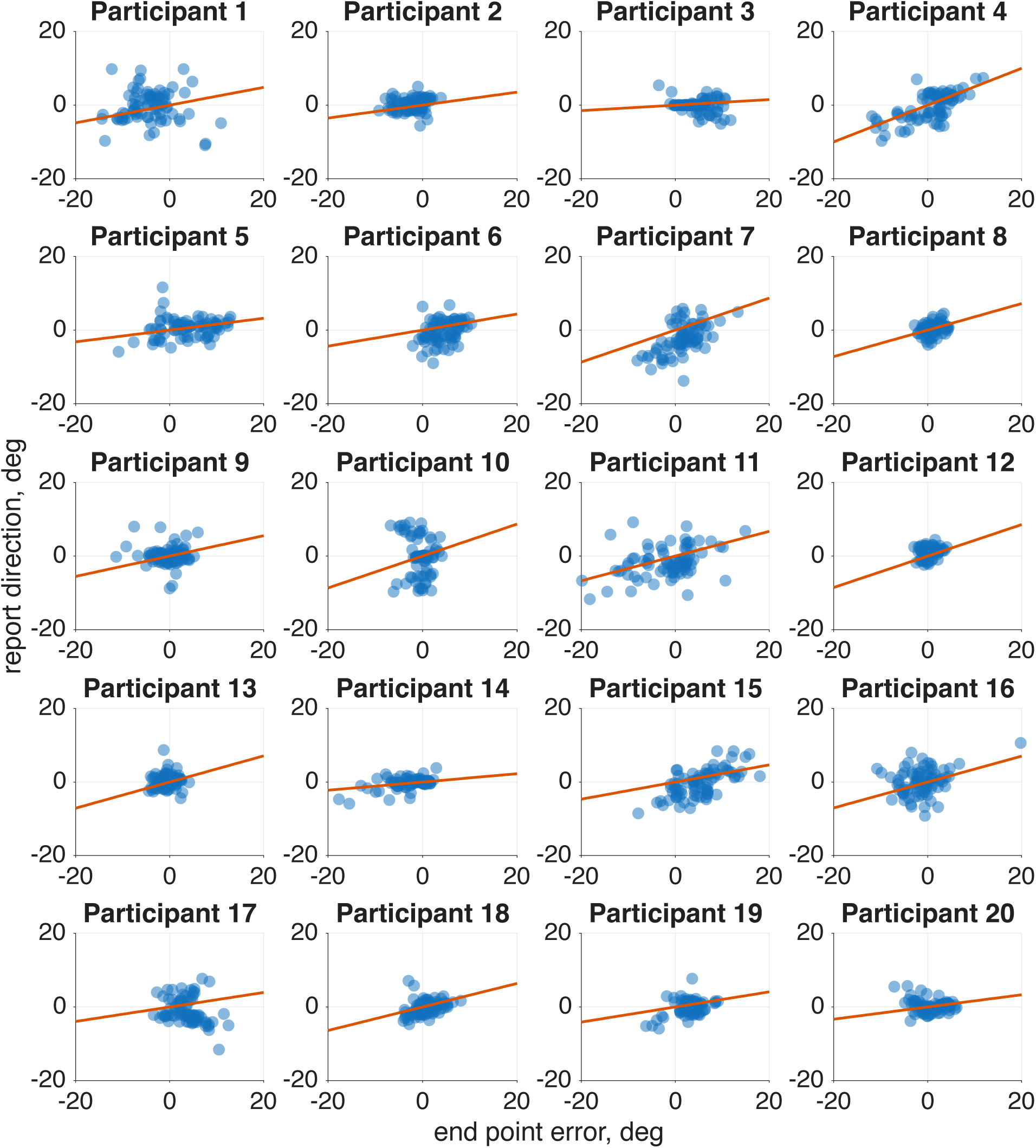
Motor awareness model fit. For all twenty participants the end point error and reported reach direction were plotted and overlaid with a line following the slope of the reliabilities of the parameters fit by the model. We can see that our model captures trends within the data.

**Supplement 2:**
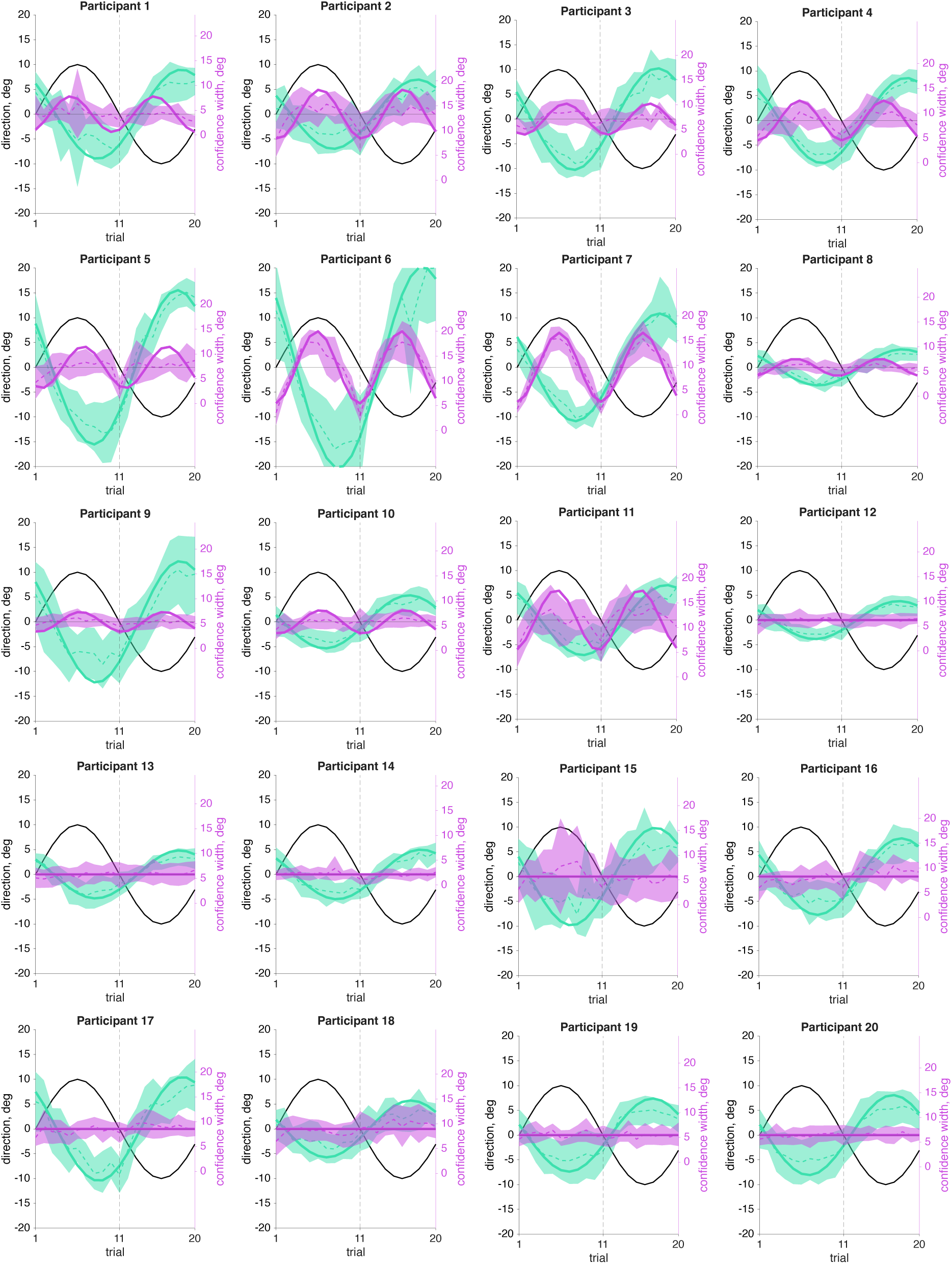
Sensorimotor confidence averaged time courses. An averaged cycle (dashed line with shaded CI) and an overlayed sine wave (solid line) using the phase and amplitude components from a Fourier analysis at the expected frequency of the response is shown here for all twenty participants. The sine wave has phase and amplitude from a Fourier analysis at the frequency of 24 cycles per session when that amplitude was significantly different from zero (Participants 1-11). For the other participants the mean is plotted (i.e., 0 amplitude). All participants showed a significant 12 cycles per session frequency for reach direction (teal).

**Supplement 3:**
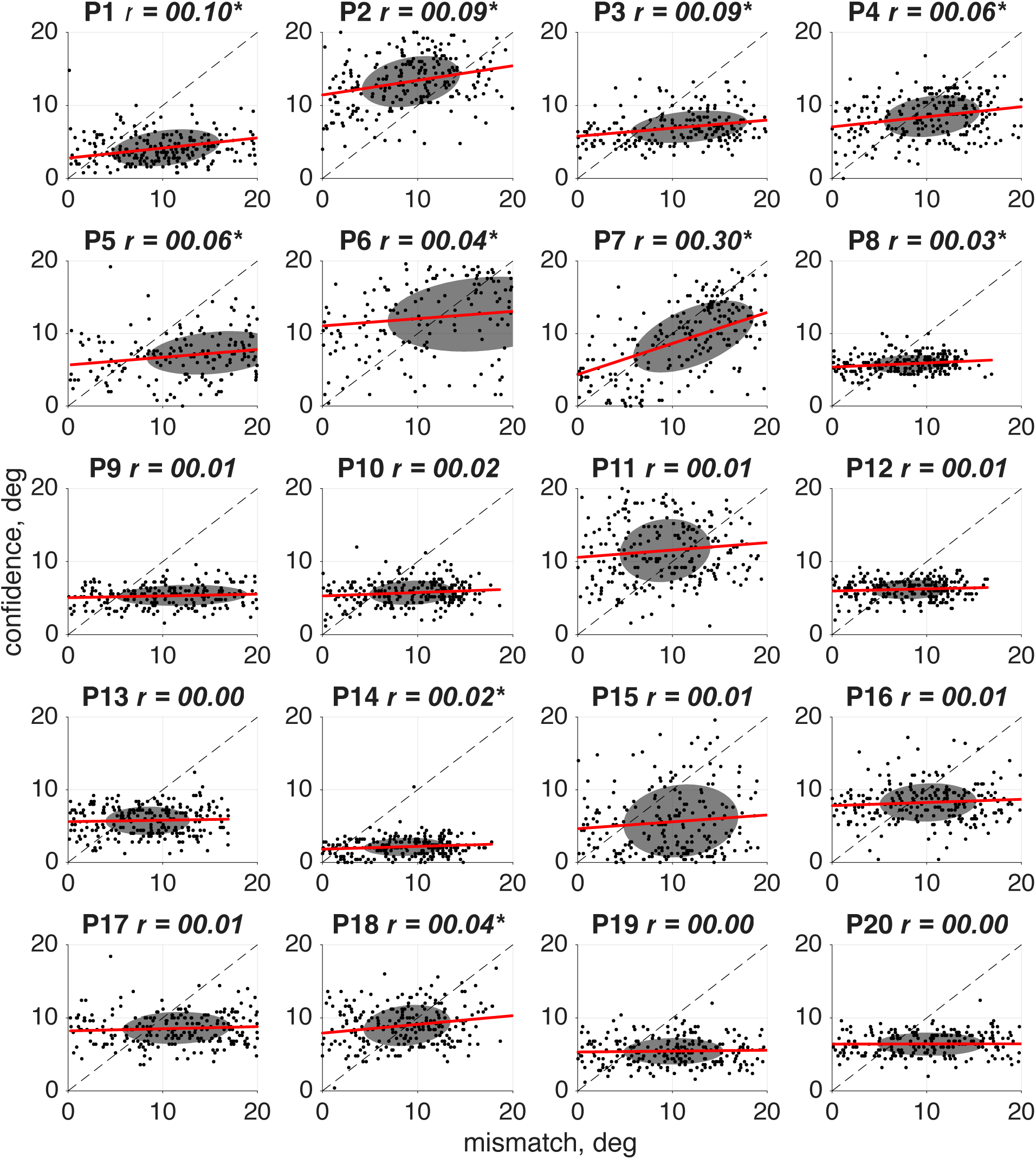
Correlation between sensorimotor confidence and the mismatch between hand and feedback position. Correlation between the confidence report on a given trial and the distance (in degrees) between the visual feedback location and hand position for the same four sample participants. Since the reach lag is on average 1.89 trials past true anti-phase, the point at which the mismatch is the smallest occurs when the error clamp is slightly off center from the target.

**Supplement 4:**
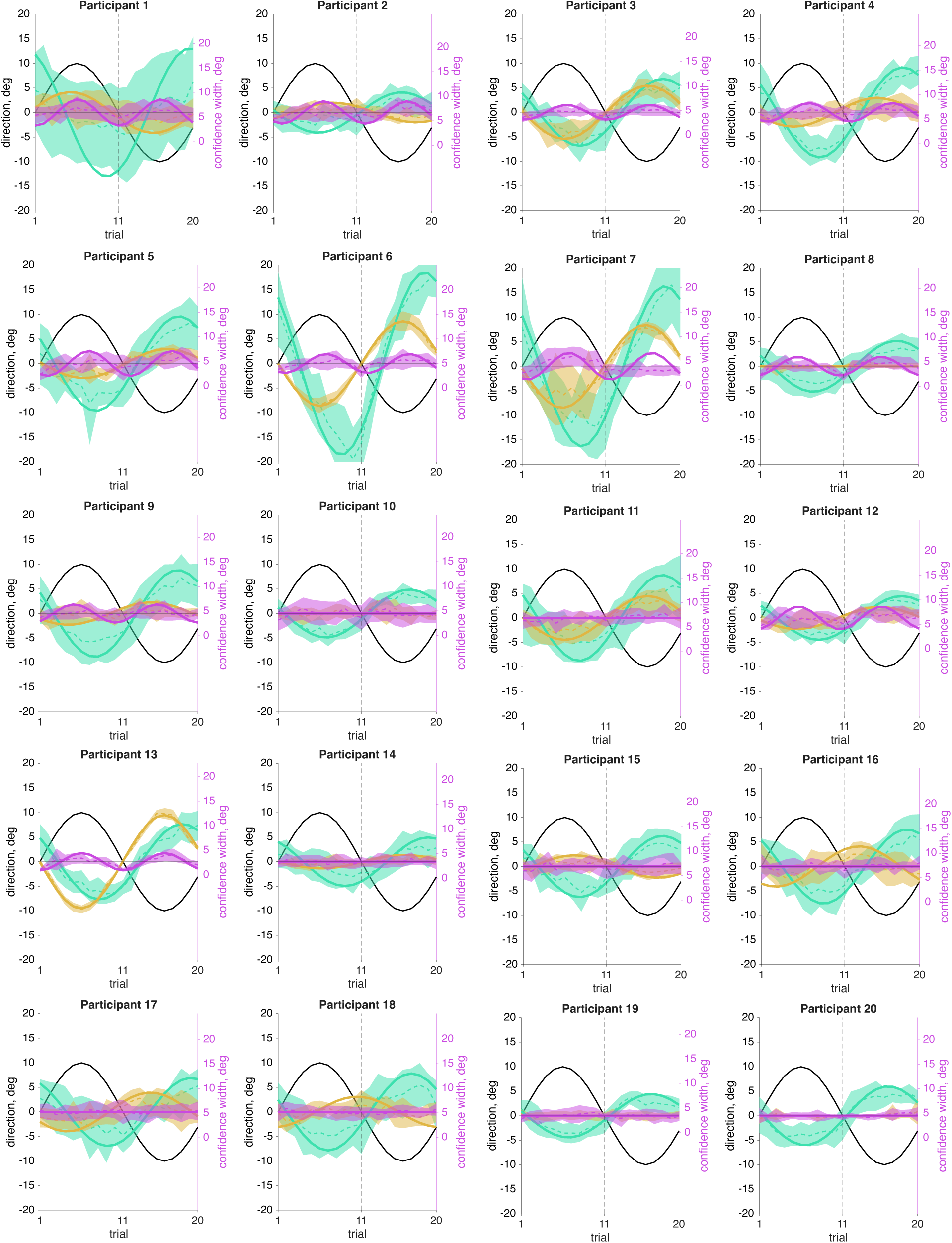
Motor-awareness confidence averaged time courses. An averaged cycle (dashed line with shaded CI) and an overlayed sine wave (solid line) using the phase and amplitude components from a Fourier analysis at the expected frequency of the response is shown here for the same four sample participants. Participants 1-9 & 12-13 had a significant 24 cycles per session frequency in confidence (magenta), while the other participants did not show this significant frequency component (and so the sine is plotted at zero amplitude). All participants showed a significant 12 cycles per session frequency for reach direction (teal). Participants 19 & 20 do not show a significant 12 cycles per session frequency in the reported hand position (tan), but the other participants all show notably different trial lags of the report relative to the error clamp.

**Supplement 5:**
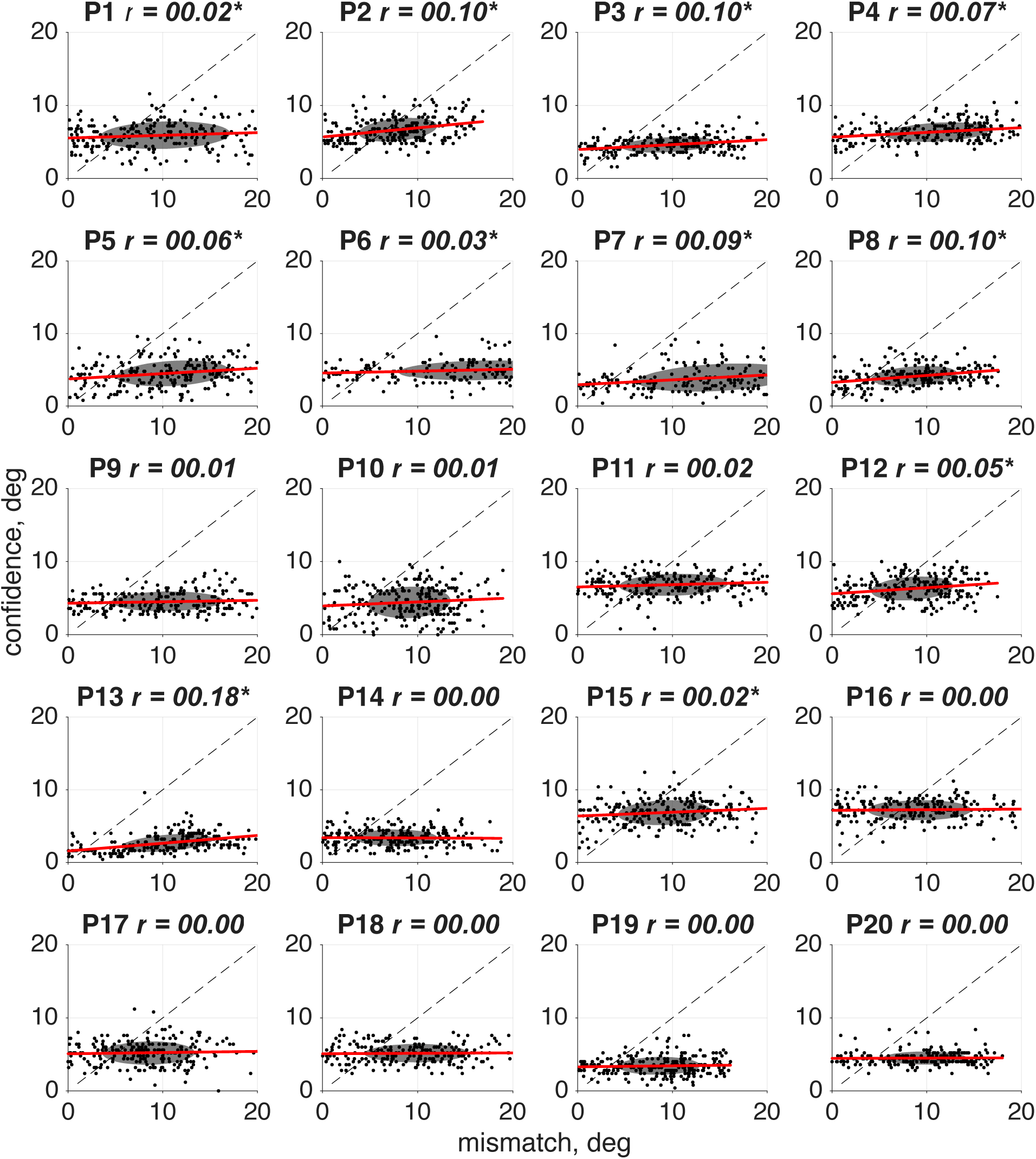
Correlation between motor-awareness confidence and the mismatch between hand and feedback position. Correlation between the confidence report on a given trial and the distance (in degrees) between the visual feedback location and hand position for the same four sample participants. Since the reach lag is on average 1.84 trials past true anti-phase, the point at which the mismatch is the smallest is not centered at the target position, 0°.

